# Structural and kinetic analysis of the COP9-Signalosome activation and the cullin-RING ubiquitin ligase deneddylation cycle

**DOI:** 10.1101/046367

**Authors:** Ruzbeh Mosadeghi, Kurt M. Reichermeier, Martin Winkler, Anne Schreiber, Justin M. Reitsma, Yaru Zhang, Florian Stengel, Junyue Cao, Minsoo Kim, Michael J. Sweredoski, Sonja Hess, Alexander Leitner, Ruedi Aebersold, Matthias Peter, Raymond J. Deshaies, Radoslav I. Enchev

## Abstract

The COP9-Signalosome (CSN) regulates cullin–RING ubiquitin ligase (CRL) activity and assembly by cleaving Nedd8 from cullins. Free CSN is autoinhibited, and it remains unclear how it becomes activated. We combine structural and kinetic analyses to identify mechanisms that contribute to CSN activation and Nedd8 deconjugation. Both CSN and neddylated substrate undergo large conformational changes upon binding, with important roles played by the N-terminal domains of Csn2 and Csn4 and the RING domain of Rbx1 in enabling formation of a high affinity, fully active complex. The RING domain is crucial for deneddylation, and works in part through conformational changes involving insert-2 of Csn6. Nedd8 deconjugation and re-engagement of the active site zinc by the autoinhibitory Csn5 glutamate-104 diminish affinity for Cul1/Rbx1 by ~100-fold, resulting in its rapid ejection from the active site. Together, these mechanisms enable a dynamic deneddylation-disassembly cycle that promotes rapid remodeling of the cellular CRL network.

## Introduction

Cullin–RING ubiquitin ligases comprise one of the largest families of regulatory enzymes in eukaryotic cells (Deshaies and Joazeiro, 2009). With as many as 240 different enzyme complexes, these E3s control a broad array of biological processes (Skaar et al., 2013). CRLs comprise seven distinct cullin–RING cores, each of which interacts with its own dedicated set of adaptor–substrate receptor complexes. Although ubiquitination by CRL enzymes is often regulated by covalent modifications of the substrate that stimulate binding to the substrate receptor, the CRL enzymes themselves are also subject to regulation.

A key mechanism that controls the activity of all known CRLs is the conjugation of the ubiquitin-like protein Nedd8 to a conserved lysine residue in the cullin subunit (e.g. K720 in human Cul1) (Enchev et al., 2015). The available structural and biochemical data indicate that Nedd8 conjugation (neddylation) stabilizes a profound conformational change in the C-terminal domain of the cullin. It loosens the interaction of the WHB domain with the RING subunit, allowing both of them to sample a greater conformational space (Duda et al., 2008), thereby enhancing the ability of the RING domain to promote ubiquitin transfer to substrate (Duda et al., 2008; Saha and Deshaies, 2008; Yamoah et al., 2008).

In addition to direct effects on ubiquitin ligase activity, Nedd8 also protects Skp1/Cul1/F-box (SCF) complexes from the substrate receptor exchange factor (SREF) Cand1 (Pierce et al., 2013; Schmidt et al., 2009; Wu et al., 2013; Zemla et al., 2013). Cand1 binds unmodified SCF complexes and promotes rapid dissociation of the F-box protein (FBP)/Skp1 substrate receptor– adaptor module from the Cul1/Rbx1 core. Cand1 can subsequently be dissociated from Cul1 by a different FBP/Skp1 complex, and as a result Cand1 functions as an SREF that accelerates the rate at which Cul1/Rbx1 comes to equilibrium with different FBP/Skp1 substrate receptor–adaptor complexes (Pierce et al., 2013). Importantly, the SREF activity of Cand1 is tightly restricted by Nedd8. Cand1 is not able to bind stably to Cul1 and promote dissociation of FBP/Skp1 when Cul1 is conjugated to Nedd8 (Liu et al., 2002; Pierce et al., 2013). These observations underscore the importance of neddylation not only for controlling the enzymatic activity of CRLs, but also potentially for controlling the repertoire of assembled CRLs.

The key role of Nedd8 in CRL biology highlights the importance of the enzymatic pathways that attach and remove Nedd8 (Enchev et al., 2015). Of particular significance is the rate of Nedd8 deconjugation, because it serves as the gateway for the exchange cycle; once Nedd8 is removed, a CRL complex is susceptible to the potent SREF activity of Cand1, and its substrate receptor can be exchanged (Pierce et al., 2013). Deconjugation of Nedd8 is mediated by the COP9-signalosome (CSN), which is an eight-subunit Nedd8 isopeptidase (Lyapina et al., 2001). The enzymatic activity of CSN resides in its Csn5 subunit, which contains a metalloprotease active site referred to as the ‘JAMM’ domain (Cope et al., 2002). The JAMM domain has the general structure E76-Xn-H138-X-H140-X10-D151 (the subscripts refer to the sequence position of these residues in human Csn5), wherein the H and D residues coordinate a zinc ion. The fourth zinc-coordination site is occupied by a water molecule that that also forms a hydrogen bond to E76 (Ambroggio et al., 2004; Sato et al., 2008; Tran et al., 2003). Deneddylation of CRLs by CSN is rapid but can be regulated by CRL substrates (Emberley et al., 2012; Enchev et al., 2012; Fischer et al., 2011). Structural analysis suggests that a CRL ubiquitination substrate bound to a substrate receptor sterically prevents concurrent binding of CSN (Enchev et al., 2012; Fischer et al., 2011). This suggests a model wherein a CRL complex has a higher probability of being conjugated to Nedd8 (and therefore of being shielded from Cand1) as long as it is bound to substrate. Upon dissociation of substrate, a race ensues between binding of either a new substrate or CSN. If CSN wins, Nedd8 is removed, paving the way for Cand1 to initiate substrate receptor exchange.

Recently, a crystal structure of free CSN was determined (Lingaraju et al., 2014). A major insight to emerge from the structure was the unexpected finding that Csn5 was present in an autoinhibited state, wherein a glutamate (Csn5-E104) within the ‘insert-1’ (INS1) sequence common to JAMM family members (Sato et al., 2008) forms a fourth ligand to the zinc, displacing the catalytic Csn5-E76-bound water molecule and shifting Csn5-E76. Csn5-E104 is found in all Csn5 orthologs, but not in other JAMM proteins, suggesting that this mode of regulation is conserved but unique to CSN. Comparison of the structure of free CSN to the structure of a catalytically-dead mutant CSN bound to Nedd8-conjugated SCF^Skp2^ determined by negative stain electron microscopy (Enchev et al., 2012) implied that binding of substrate to CSN may induce several conformational changes in the latter, including movement of the N-terminal domains (NTD) of Csn2 and Csn4 towards the cullin. The latter movement, in turn, might be further propagated to the Csn5/6 module (Lingaraju et al., 2014). Moreover, it is reasonable to expect that during catalysis INS1 moves out of the active site and Csn5-E76 adopts a position similar to that observed in a crystallographic structure of Csn5 in isolation (Echalier et al., 2013). Interestingly, if Csn5-E104 is mutated to an alanine, CSN more rapidly cleaves the simple model substrate ubiquitin-rhodamine (Lingaraju et al., 2014). This was interpreted to mean that the primary reason for the autoinhibited state is to keep CSN off until it binds a physiologic substrate, which would prevent spurious cleavage of non-cullin Nedd8 conjugates and possibly even ubiquitin conjugates. However, the full extent of the conformational changes required to form an activated complex between CSN and its neddylated substrate, as well as the detailed molecular basis for these changes remain to be established. Therefore, at present, the mechanism of how CSN is switched on and off and the significance of this switching behavior remain unknown.

## Results

### Structural insights from cryo EM and single particle analysis of a CSN-SCF-Nedd8^Skp2/Cks1^ complex

To gain detailed insights into the molecular determinants underlying activation of CSN, we performed cryo electron microscopy (cryo EM) and single particle analysis of CSN^5H138A^ (we use the nomenclature CSN^#x^ where # refers to subunit number and x to the specific mutation) in complex with neddylated SCF^Skp2/Cks1^ (the sample is described in (Enchev et al., 2012) (Figure 1A, Figure 1–figure supplements 1A–D). The Csn5-H138A mutant lacks one of the JAMM ligands that coordinate the catalytic zinc. This mutant forms a normal CSN complex that has been extensively characterized (Enchev et al., 2012). We used ~75000 single molecular images for the final three-dimensional reconstruction and the structure was refined to a nominal resolution of 7.2 Å, according to the ‘gold standard’ criterion of a Fourier shell correlation (FSC) of 0.143 (Rosenthal and Henderson, 2003; Scheres and Chen, 2012) (Figure 1–figure supplements 1C–D). However, some regions in the density map were better defined than others (see below). To avoid over-interpretation, for the subsequent analysis we low-pass filtered the map to 8.5 Å, according to the more stringent criterion of an FSC of 0.5.

**Figure 1.**
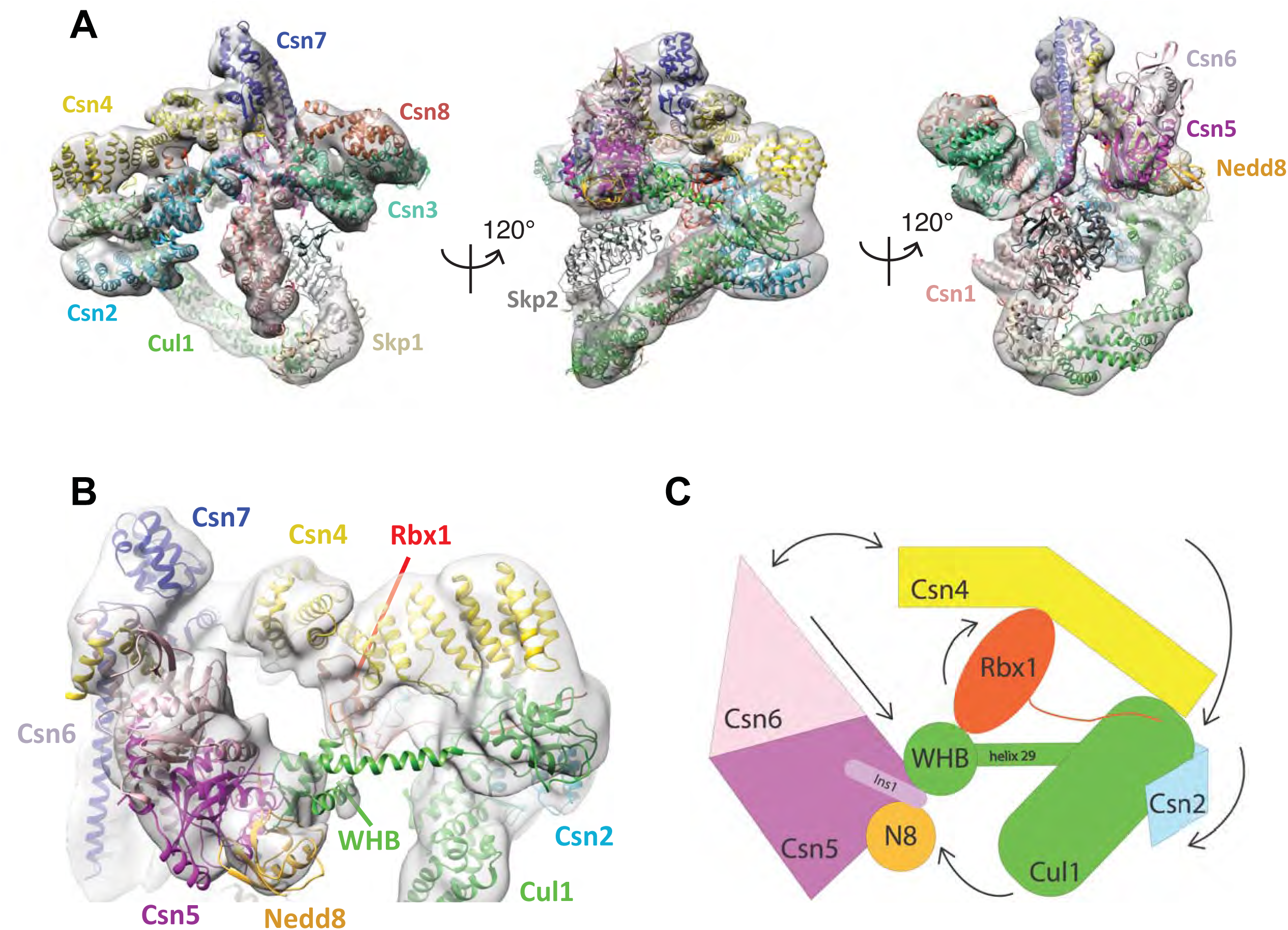
**Cryo-electron microscopy of a CSN-SCF complex**. (A) Molecular model of CSN^5H138A^-SCF-N8^Skp2/Cks1^ docked into the cryo-electron density map (gray mesh). (B) Close-up view of the model, showing the observed conformations of Csn2, Csn4, Rbx1, Csn5/6 and WHB-Nedd8 and (C) a cartoon representation of the differences between the apo CSN and substrate-bound state.

The cryo EM structure reported here, alongside the available crystal structure of CSN (Lingaraju et al., 2014), enabled us to visualize a broad array of conformational changes that take place upon complex formation in both CSN and neddylated Cul1/Rbx1, well beyond what was possible with the prior lower resolution model based on negative stain EM (Figure 1). Specifically, this allowed us to describe movements of the N-terminal domains of Csn2 and Csn4, the MPN domains of Csn5 and Csn6. Moreover, in contrast to our previous work, we could locate the RING domain of Rbx1, as well as Nedd8 and the winged-helix B (WHB) domain of Cul1 relative to Csn5. Nevertheless, the present resolution precludes the determination of the exact orientations of the latter domains but notably, the relative positions of the RING, WHB and Nedd8 reported here have not been reported in any structural model of a cullin, and strongly suggest that both the enzyme and substrate undergo significant conformational rearrangements to enable catalysis.

To obtain the model shown in Figure 1, we initially docked the crystal structure of CSN (Lingaraju et al., 2014) and a model of Cul1-Nedd8/Rbx1/Skp1/Skp2/Cks1 (Enchev et al., 2012) as rigid bodies into the electron density map (Figure 1–figure supplements 1E–H). We observed very good matches between the respective map segments and the atomic coordinates for the scaffold subunits Csn1, Csn3, Csn7 and Csn8, the winged-helix domains of Csn2 and Csn4 (Figure 1–figure supplement 1E), and the helical bundle formed by the C-termini of all eight CSN subunits (Figure 1–figure supplement 1F) as well as the expected recovery of secondary structure at this resolution. Similarly, there was a very good overlap between the coordinates of Cul1 (with the exception of helix29 and the WHB domain, see below) and Skp1 and the corresponding electron density segments (Figure 1–figure supplement 1G). However, the local resolution was lower without recovery of secondary structure in the N-terminal domain of Cul1. Moreover, the density of the substrate receptor Skp2/Cks1 was poorly defined (Figure 1–figure supplement 1G), indicating a potential flexibility in this region. Since the presence of Skp1/Skp2 had modest effects on the affinity and deneddylation activity (see below), we did not interpret this observation further.

In contrast to the large segments of CSN that were unaltered upon binding substrate, there was nearly no overlap between the EM density map and the N-terminal portions of Csn2 and Csn4, as well as the MPN-domains of Csn5 and Csn6, the RING domain of Rbx1, the WHB domain of Cul1, and Nedd8 (Figure 1–figure supplement 1H). We thus docked these domains individually (Figure 1B, Movie 1). A Csn2 N-terminal fragment encompassing the portion between its crystallographically resolved N-terminus (amino acid 30) through to a flexible loop at amino acid 180, was docked as a rigid body (Figure 1–figure supplement 1I), positioning it close to the four-helical bundle and helix 24 of Cul1 (Zheng et al., 2002). An N-terminal fragment of Csn4, spanning amino acids 1 to 295, which ends in a previously reported hinge loop (Lingaraju et al., 2014), was also docked independently as a rigid body (Figure 1–figure supplement 1J). The resulting conformation of Csn4 resembles a crystal form of Csn4 observed in isolation (Lingaraju et al., 2014). The two N-terminal helical repeat motifs of Csn4 make contacts with the winged-helix A domain of Cul1 (Figure 1B and Figure 1–figure supplement 1J, right hand panel, red arrow and green circle). Moreover, these positions of Csn2 and Csn4 delineated a density in the map, which could accommodate the RING domain of Rbx1 (Figure 1B and Figure 1–figure supplement 1J, right hand panel, black ellipse), with the RING proximal to two conserved helices between amino acids 160 and 197 of Csn4 (Figures 1B and Figure 1–figure supplement 1K, black arrow) and a loop in Csn2 located between residues 289 and 306. The exact orientation of the RING domain awaits a structure at higher resolution.

To improve the fit of Csn5 and Csn6, we moved their MPN domains as rigid bodies into the neighboring map segment of similar shape and dimensions (Figure 1–figure supplement 1L). The local resolution in this region was lower, presumably due to higher flexibility around the catalytic site. Importantly, after docking Csn5, we observed two empty neighboring densities (Figure 1– figure supplement 1L, right hand panel, circles), which accommodated the two yet undocked protein components – Nedd8 and the WHB domain of Cul1 (Figure 1–figure supplement 1M). The docking of the latter was enabled by allowing helix 29 and the WHB, amino acids 690 to the C-terminus of Cul1, to move as a rigid body towards Csn5. However, we did not observe an electron density around helix 29 of Cul1, consistent with a structural flexibility in this region. This model places the neddylated WHB domain in close binding proximity to the RING domain, as well as both INS1 and INS2 of Csn5. The hydrophobic patch of Nedd8 is facing INS1, and not the WHB domain, as has been reported for the isolated neddylated C-terminal domain of Cul5 (Duda et al 2008). Similar to the RING domain, we cannot be fully certain about the exact orientations of Nedd8 and the WHB domain at the present resolution. Nevertheless, to further substantiate this docking, we mutagenized conserved charged residues in the INS2 domain of Csn5 as well as the WHB domain, and as expected all of these constructs showed reduced catalytic activity in deneddylation assays (Figure 1–figure supplement 1N, O).

We sought orthogonal experimental validation for the molecular docking of the individual subunits and domains in the electron density map by performing cross-linking coupled to mass spectrometric analysis of the cross-linked peptides (Leitner et al., 2014) following the procedure described in Birol et al. (2014) (supplementary files 1-6). For the cross-linker used in this study (disuccinimidylsuberate H_12/_D_12_), the maximum predicted distance between two cross-linked lysine residues is generally accepted to be below ~ 30 Å (Politis et al., 2014). As shown in supplementary files 1, out of the 39 high-confidence inter-subunit cross-links detected within the CSN^5H138A^–N8-SCF^Skp2/Cks1^ complex at a false discovery rate (FDR) of 5 percent, the great majority was within regions of modeled atomic structure and only six links exhibited a distance larger than 30 Å when mapped onto our model. However, all of these larger-distance links are connected to the flexibly positioned Skp2 density. Moreover, we further performed similar cross-linking experiments on a number of different CSN-CRL complexes, varying the substrate receptor, the cullin and the neddylation state (supplementary files 2-6). All results were consistent with the architecture proposed here for CSN^5H138A^-N8-SCF^Skp2/Cks1^. Intriguingly, when taking into account cross-links with an FDR of up to 0.25 (supplementary files 2 and 3), we found two cross-links that support proximity of K290 in Csn4 and K89 in the RING domain (supplementary file 2), as well as K32 in Csn4-NTD and K587 in Cul1, which is in the immediate vicinity of the WHA domain of Cul1 (supplementary file 3), as suggested by our EM reconstruction.

### Development and validation of an assay to measure binding of CSN to substrate and product

To understand how the structure of CSN and the CSN–SCF complex relates to substrate binding and the mechanism of deneddylation, we sought to develop quantitative binding assays to measure interaction of CSN with its substrates and products. To this end, the environmentally-sensitive dye dansyl was conjugated to the C-terminus of Cul1 using ‘sortagging’ (Theile et al., 2013) to generate dansylated Cul1/Rbx1 (Cul1^d^/Rbx1) (Figure 2–figure supplement 2A). Cul1^d^/Rbx1 exhibited normal E3 activity (Figure 2–figure supplement 2B) and bound CSN with an affinity similar to Cul1/Rbx1 based on their IC_50_ values for competitive inhibition of a deneddylation reaction (Figure 2–figure supplement 2C; Emberley et al., 2012). When Cul1^d^/Rbx1 was incubated with CSN (all CSN preparations used in this work are shown in Figure 2–figure supplement 2D), we observed an increase in dansyl fluorescence (Figure 2A). This signal was due to specific binding because it was chased upon addition of excess unlabeled Cul1/Rbx1 (Figure 2A, titration shown in Figure 2–figure supplement 2E) or Cand1 (Figure 2–figure supplement 2F), which competes for substrate deneddylation by CSN (Emberley et al., 2012; Enchev et al., 2012). Thus, we concluded that the increase in dansyl fluorescence accurately reported on the interaction of CSN with Cul1^d^/Rbx1. Using this assay we determined that CSN bound Cul1^d^/Rbx1 with a *K_d_* of 310 nM (Figure 2B). Cul1^d^/Rbx1 binding to CSN was only modestly affected by the addition of free Nedd8 (Figure 2–figure supplement 2G) or assembly with Skp2/Skp1 (Figure 2–figure supplement 2H) or Fbxw7/Skp1 (Figure 2–figure supplement 2I).

**Figure 2.**
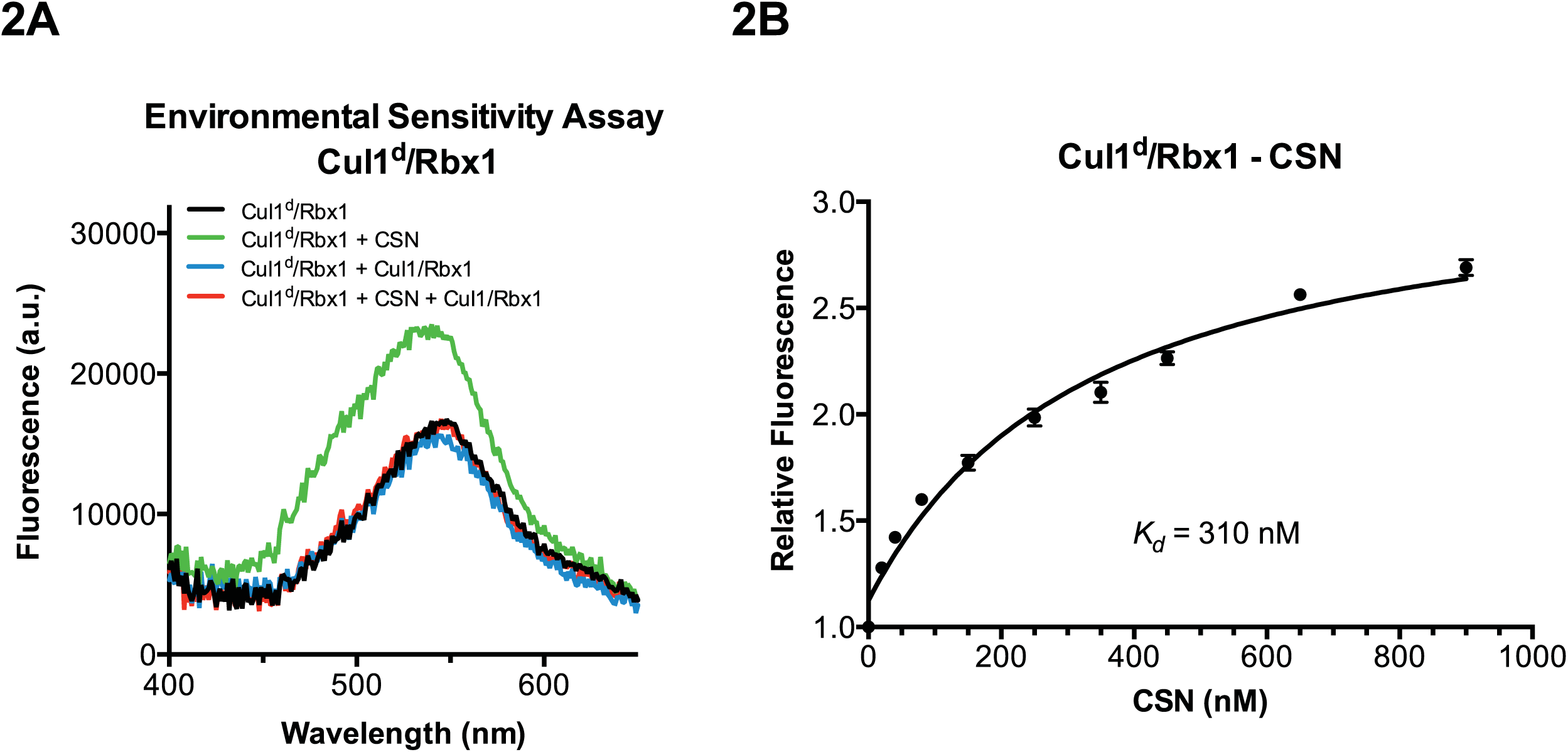
**Development and validation of a binding assay for CSN–Cul1/Rbx1 interaction**. (A) Equilibrium binding of CSN to Cul1^d^/Rbx1 and competition by unlabeled Cul1/Rbx1. The indicated proteins were mixed and allowed to equilibrate prior to determination of dansyl fluorescence in a fluorometer. Note that Cul1/Rbx1 blocks the fluorescence enhancement caused by CSN. CSN, Cul1^d^/Rbx1, and Cul1/Rbx1 were used at 350, 30, and 4000 nM, respectively. (B) Equilibrium binding of CSN to Cul1^d^/Rbx1. Cul1^d^/Rbx1 (30 nM) was mixed with increasing concentrations of CSN and the proteins were allowed to equilibrate prior to determining the change in dansyl fluorescence in triplicate samples. Error bars represent standard deviation.

We next sought to measure binding of neddylated Cul1^d^/Rbx1 (Cul1^d^-N8/Rbx1) to CSN but it was not possible because the substrate was rapidly deneddylated. To circumvent this problem, we performed binding assays with the extensively characterized inactive mutant CSN^5H138A^ (assay confirming loss of activity is shown in Figure 3–figure supplement 3A). Remarkably, CSN^5H138A^ bound Cul1^d^-N8/Rbx1 ~200-fold more tightly than CSN bound Cul1^d^/Rbx1 (*K_d_* 1.6 nM vs. 310 nM; Figures 3A–B). Note that the estimated *K_d_* falls well below the fixed concentration of Cul1^d^-N8/Rbx1 used in the assay. This introduces greater uncertainty into our estimate but nevertheless we can conclude with confidence that the binding of substrate to CSN^5H138A^ is very tight (≤5 nM; see Materials and Methods for further discussion of this matter). As reported above for CSN binding to product, addition of Skp2/Skp1 or Fbxw7/Skp1 had comparatively minor effects on affinity (Figure 3–figure supplements 3B–C). Thus, for the sake of simplicity, we used Cul1^d^/Rbx1 heterodimer for the remaining binding experiments.

**Figure 3.**
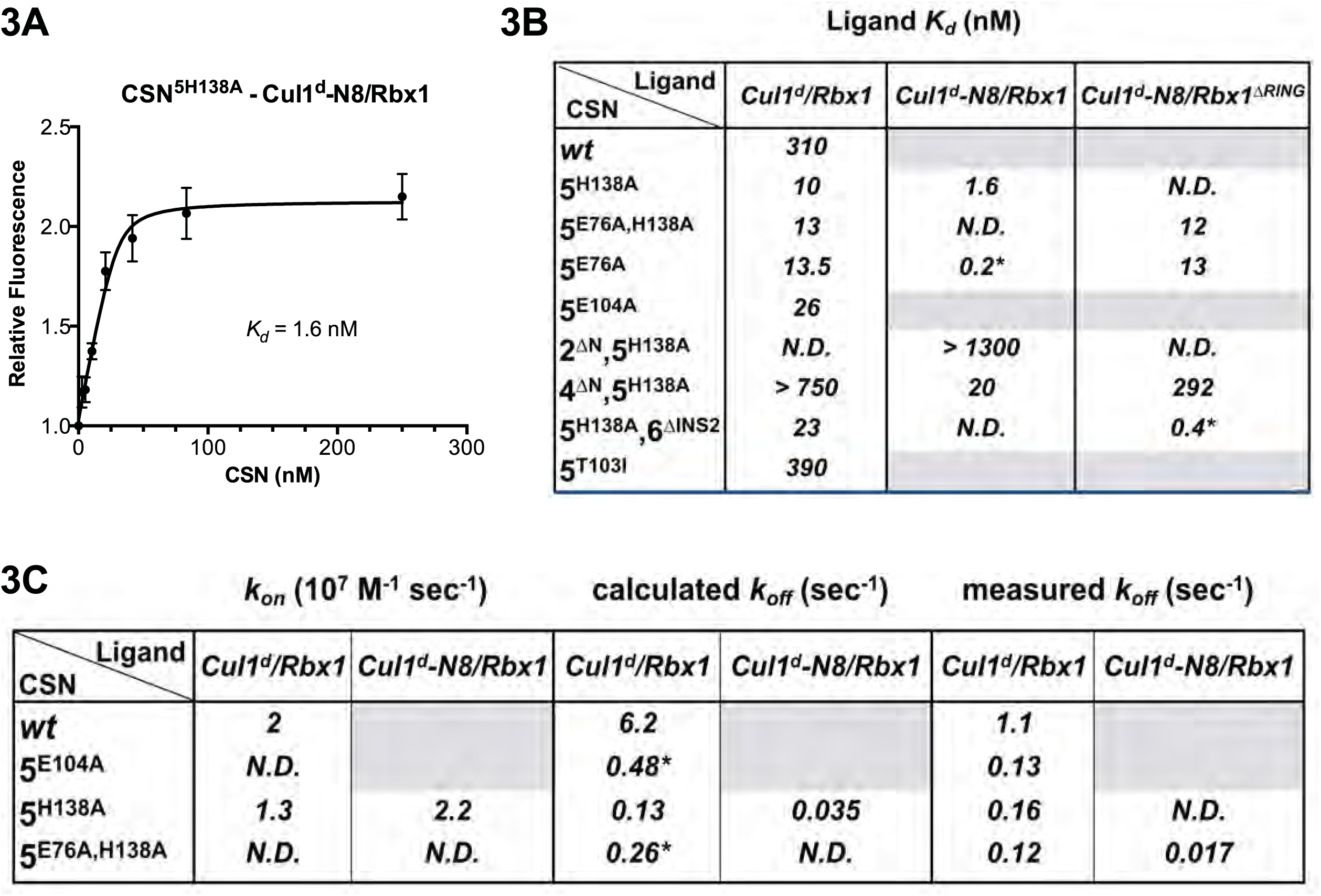
**Quantitative determination of enzyme–substrate binding affinities for wild type and mutant proteins**. (A) Tight binding of CSN^5H138A^ to substrate. Cul1^d^-N8/Rbx1 and CSN^5H138A^ were mixed and allowed to equilibrate prior to determining the change in dansyl fluorescence. (B) Summary of *K_d_* measurements for the indicated CSN complexes tested against unmodified Cul1^d^/Rbx1, Nedd8-conjugated Cul1^d^-N8/Rbx1 or Cul1^d^-N8/Rbx1^ΔRING^ ligand. Boxes shaded in gray indicate combinations that could not be analyzed due to deneddylation during the binding reaction. For some complexes that bound weakly it was not feasible to titrate to saturation and so a lower boundary for *K_d_* is indicated. N.D., not determined. *, due to the configuration of our assay, extremely low *K_d_* values cannot be reliably determined. (C) Summary of *k_on_* and *k_off_* measurements for the indicated CSN complexes tested against Cul1^d^/Rbx1 or Cul1^d^-N8/Rbx1. Each reported *k_off_* is the mean of at least 8 replicates. For comparison, *k_off_* values calculated from *k_on_* and *K_d_* measurements are also shown. For cases where *k_on_* was not measured (marked with asterisks), we assumed a value that was the average (1.83 x 107 M-1 sec^-1^) of the three measured *k_on_* values. Boxes shaded in gray indicate combinations that could not be analyzed due to deneddylation upon complex formation. N.D., not determined.

The strikingly high affinity we observed for binding of CSN^5H138A^ to Cul1^d^-N8/Rbx1 led us to question whether it was mainly due to Nedd8 or whether the H138A mutation might also enhance affinity. To this end, we measured binding of CSN^5H138A^ to Cul1^d^/Rbx1 and observed an unexpectedly low *K_d_* of ~10 nM (Figure 3B, Figure 3–figure supplement 3D), which was confirmed with an independent preparation of CSN^5H138A^ (Figure 3–figure supplement 3E). Thus, neddylation improved affinity of Cul1^d^/Rbx1 for CSN^5H138A^ by ~6-fold, whereas the Csn5-H138A mutation improved affinity for Cul1^d^/Rbx1 by ~30-fold. The high affinity binding of CSN^5H138A^ to substrate was supported by an orthogonal competition experiment in which 100 nM CSN^5H138A^ completely blocked deneddylation of 75 nM Cul1-N8/Rbx1 (Figure 3–figure supplement 3A). We considered the possibility that the Csn5-H138A mutation might enable formation of an aberrant, super-tight enzyme:substrate ([ES]) complex that does not normally form between the wild type proteins. However, as will be described later on, this hypothesis was rejected based on kinetic arguments.

We next sought to determine whether the large differences we observed in *K_d_* values were due to differences in *k_on_* or *k_off_*. Remarkably, despite a 200-fold difference in *K_d_* for CSN^5H138A^ binding to substrate compared to CSN binding to product, the *k_on_* values for formation of these complexes were nearly identical (2.0 x 10^7^ M^-1^ sec^-1^ for CSN–product and 2.2 x 10^7^ M^-1^ sec^-1^ for CSN^5H138A^–substrate; Figure 3C). This suggested that the difference in affinity was driven by a large difference in *k_off_*. To test this hypothesis, we directly measured *k_off_* values for select [ES] and enzyme-product complexes by pre-forming the complex and then adding excess unlabeled Cul1/Rbx1 chase and monitoring the reduction in dansyl fluorescence over time (Figure 3C and Figure 3–figure supplement 3F-I; for this and a subsequent experiment in Figure 4B, we used CSN^5E76A/5H138A^ in one of the assays instead of CSN^5H138A^; the double mutant behaved like CSN^5H138A^ in that it bound Cul1^d^/Rbx1 with the same affinity as shown in Figure 3–figure supplement 3J.). Consistent with the predictions from the *K_d_* and *k_on_* values, substrate dissociated very slowly from CSN^5E76A,5H138A^, whereas product dissociated ~65-fold faster from CSN. This suggests that as substrate is deneddylated to product, its affinity for CSN is strongly reduced and its *k_off_* speeds up.

**Figure 4.**
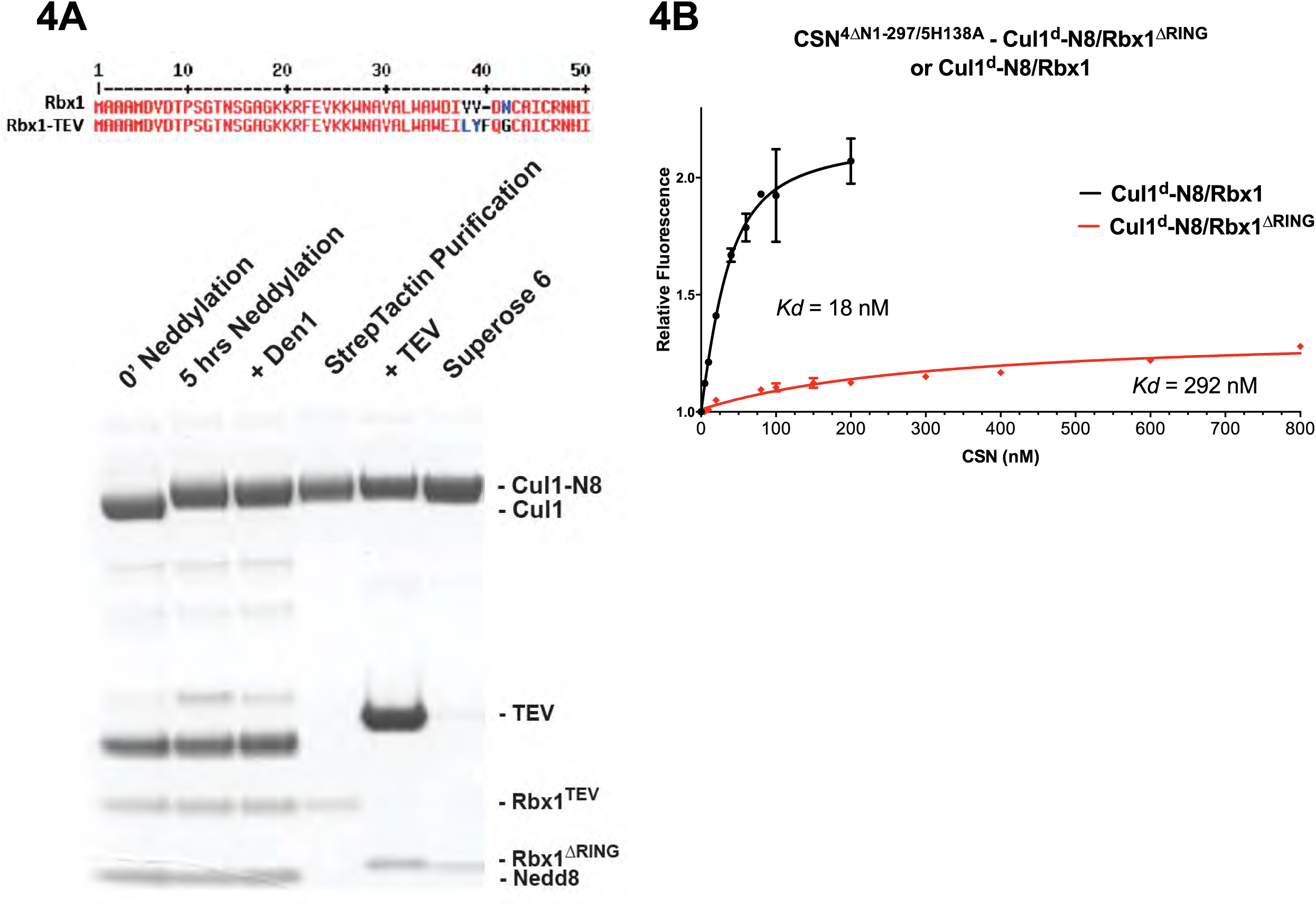
**The N-terminal domains of Csn2 and Csn4 and the RING domain of Rbx1 play key roles in substrate binding and deneddylation**. (A) Generation of Cul1-N8/Rbx1^ΔRING^. Top: a TEV protease site was engineered between the N-terminal β-strand and the RING domain of Rbx1 as indicated. Only the first 50 amino acids of Rbx1 are shown. Bottom: Purified protein was subjected to the indicated treatments (see Materials and Methods for details) and reactions were fractionated by SDS-PAGE and stained with Coomassie Blue. (B) Deletion of the Csn4-NTD and Rbx1-RING domains independently reduce affinity of CSN for substrate. The indicated proteins were mixed and allowed to equilibrate prior to determining the change in dansyl fluorescence in triplicate samples. Error bars represent standard deviation. *K_d_* values measured in this experiment are also reported in Figure 3B.

### The N-terminal domains of Csn2 and Csn4 and the RING of Rbx1 promote enzyme–substrate interaction

Armed with assays to measure binding and deneddylation of substrate, we next sought to test the implications that emerged from our structural analysis of the CSN^5H138A^–N8-SCF^Skp2/Cks1^ complex. First, we investigated the roles of the NTDs of Csn2 and Csn4, both of which, upon binding substrate, underwent conformational changes and made contact with Cul1 and the RING domain of Rbx1 (Figures 1B–C, Figure 1–figure supplements 1I–K) (Lingaraju et al., 2014). To measure the effect of these mutations on binding to Cul1^d^-N8/Rbx1, we combined them with Csn5-H138A to prevent deneddylation. Deletion of the first 269 amino acids of Csn2, observed to interact with Cul1 but not the RING domain of Rbx1, caused a massive loss in binding to substrate (*K_d_* > 1300 nM; Figure 3B, Figure 3–figure supplement 3K). Thus, the contact we observed between Csn2-NTD and N8-SCF^Skp2/Cks1^ was critical to formation of the [ES] complex. By contrast, deletion of the first 297 amino acids NTD of Csn4 (4ΔN), a portion which was observed to form interfaces with both Cul1 and the RING domain of Rbx1, had a relatively modest effect; CSN^4ΔN,5H138A^ bound Cul1^d^/Rbx1 and Cul1^d^-N8/Rbx1 with *K_d_* values of > 750 nM and 20 nM, respectively (Figure 3B, Figure 3–figure supplement 3L-M).

In addition to the motions of the Csn2 and Csn4 NTDs, our structural analysis revealed formation of substantial interfaces between CSN and the RING domain of Rbx1. To test the role of the RING domain in complex formation, we generated both Cul1/Rbx1 and Cul1^d^/Rbx1 in which the RING domain can be deleted by introducing a TEV protease cleavage site (Dougherty et al., 1989) after residue 37 of Rbx1 to generate Cul1 (or Cul1^d^)/Rbx1^TEV^ (Figure 4A). This was essential, because it would not be possible to conjugate Nedd8 to Cul1/Rbx1 expressed as a mutant lacking the RING domain. After conjugating Nedd8 to the purified complex, we treated it with TEV protease to remove the RING domain, yielding Cul1 (or Cul1^d^)-N8/Rbx1^ΔRING^ (Figure 4A). The truncated Cul1/Rbx1^ΔRING^ was inactive in an ubiquitylation assay (Figure 4–figure supplement 4A) but behaved as a monodisperse sample with the expected hydrodynamic radius upon size exclusion chromatography (Figure 4–figure supplement 4B). Notably, Cul1^d^-N8/Rbx1^ΔRING^ bound CSN^5E76A,5H138A^ and CSN^5E76A^ with affinities (12 nM and 13 nM respectively; Figure 3B, Figure 3– figure supplement 3N) similar to that observed for binding of wild type Cul1^d^-N8/Rbx1 to CSN^4ΔN,5H138A^. Given the similar effects of the Csn4-ΔNTD and Rbx1-ΔRING mutations on complex formation, we next tested whether their effects arose from loss of the interface that forms between these domains (Figure 1–figure supplement 1K). However, double mutant analysis suggested that the Csn4-ΔN and Rbx1-ΔRING mutations had largely independent effects on binding (Figure 4B). The overall picture that emerged from these studies in light of the structural data is that the interaction of Csn2-NTD with neddylated substrate makes a large contribution to binding energy, with modest enhancements independently provided by the Csn4-NTD and Rbx1-RING domains.

### The ‘E-vict’ enables efficient clearance of product from the CSN active site

The striking difference in the *K_d_* for CSN^5H138A^ binding to substrate compared to CSN binding to product suggested that a conformational rearrangement of the [ES] complex occurs upon cleavage of the isopeptide bond, resulting in a large increase in the product *k_off_*, thereby preventing the enzyme from becoming product-inhibited. However, we were puzzled by the relatively minor impact of Nedd8 on the affinity of Cul1^d^/Rbx1 for CSN^5H138A^; whereas substrate bound with *K_d_* of 1.6 nM, product binding was only ~6-fold weaker (Figure 3B). Why, then, did CSN bind so much less tightly to product? We reasoned that a key difference between CSN^5H138A^ and CSN is the absence of the active site zinc from CSN^5H138A^, which prevents formation of a stable apo-CSN complex in which E104 of the INS1 domain of Csn5 is bound to the active site zinc. If this conjecture is correct, it makes the prediction that CSN^5E104A^, which should also be unable to form stable apo-CSN, should likewise exhibit high affinity for product. This was confirmed: CSN5e104 bound Cul1^d^/Rbx1 with a *K_d_* of 26 nM (Figure 3B, Figure 3–figure supplement 3O). Furthermore, measurement of *k_off_* values revealed that product dissociated from CSN^5H138A^ and CSN^5E104A^ about 8-fold more slowly than it dissociated from CSN (Figure 3C). Based on these observations, we propose the ‘E-vict’ hypothesis, which is described in more detail in the Discussion. The essence of this hypothesis is that, following cleavage of the isopeptide bond and dissociation of Nedd8, INS1 of Csn5 engages the active site zinc. This accelerates the rate of dissociation of deneddylated Cul1/Rbx1, thereby preventing CSN from becoming clogged with product. We note that Csn5-E76 also contributes to the operation of this mechanism, because CSN^5E76A^ bound tightly to product (Figure 3–figure supplement 3P). We speculate that engagement of the active site zinc by Csn5-E104 forces Csn5-E76 into a configuration that promotes egress of product. Further insights into the exact sequence of events that accelerates product dissociation await high-resolution structures of CSN bound to Cul1/Rbx1 in various states.

### Kinetic effects of binding-defective mutations on substrate deneddylation

We next sought to address the effects of the enzyme and substrate mutations described in the preceding sections on the deneddylation reaction. We previously showed that CSN2Δn has severely reduced catalytic activity (Enchev et al., 2012), which is consistent with the binding data reported here. CSN4Δn exhibited a 20-fold defect in substrate cleavage (Figure 5A, Figure 5–figure supplement 5A). Meanwhile, the *k_cat_* for cleavage of Cul1-N8/Rbx1^ΔRING^ by CSN was reduced by a staggering ~18,000-fold relative to wild type substrate (Figures 5A–B). Given that the neddylated ΔRING substrate bound to CSN with only modestly reduced affinity, we surmised that the principal defect of this mutant might be its failure either to induce the activating conformational change in CSN, and/or to position accurately the isopeptide bond in the active site. Although we do not have the tools to address the latter point, we queried the former by examining the Csn6-ΔINS2 mutation, which partially mimics the effect of substrate binding in that it destabilizes the autoinhibited state (Lingaraju et al., 2014). The Csn6-ΔINS2 mutation slightly weakened binding to wild type product (Figure 3B, Figure 3–figure supplement 3Q) but completely suppressed the modest binding defect of the neddylated ΔRING substrate (Figure 3B, Figure 3–figure supplement 3R) and promoted an ~8-fold increase in its deneddylation rate (Figure 5–figure supplement 5B). This partial suppression effected by Csn6-ΔINS2 suggests that the RING domain contributes to the constellation of conformational changes in CSN that occur upon substrate binding. Note that the CSN^6Δins2^ enzyme nevertheless exhibited a > 1,000-fold defect towards the Cul1-N8/Rbx1^ΔRING^ substrate, strongly indicating further functions of the RING domain, which may include a potential role in substrate positioning as well.

**Figure 5.**
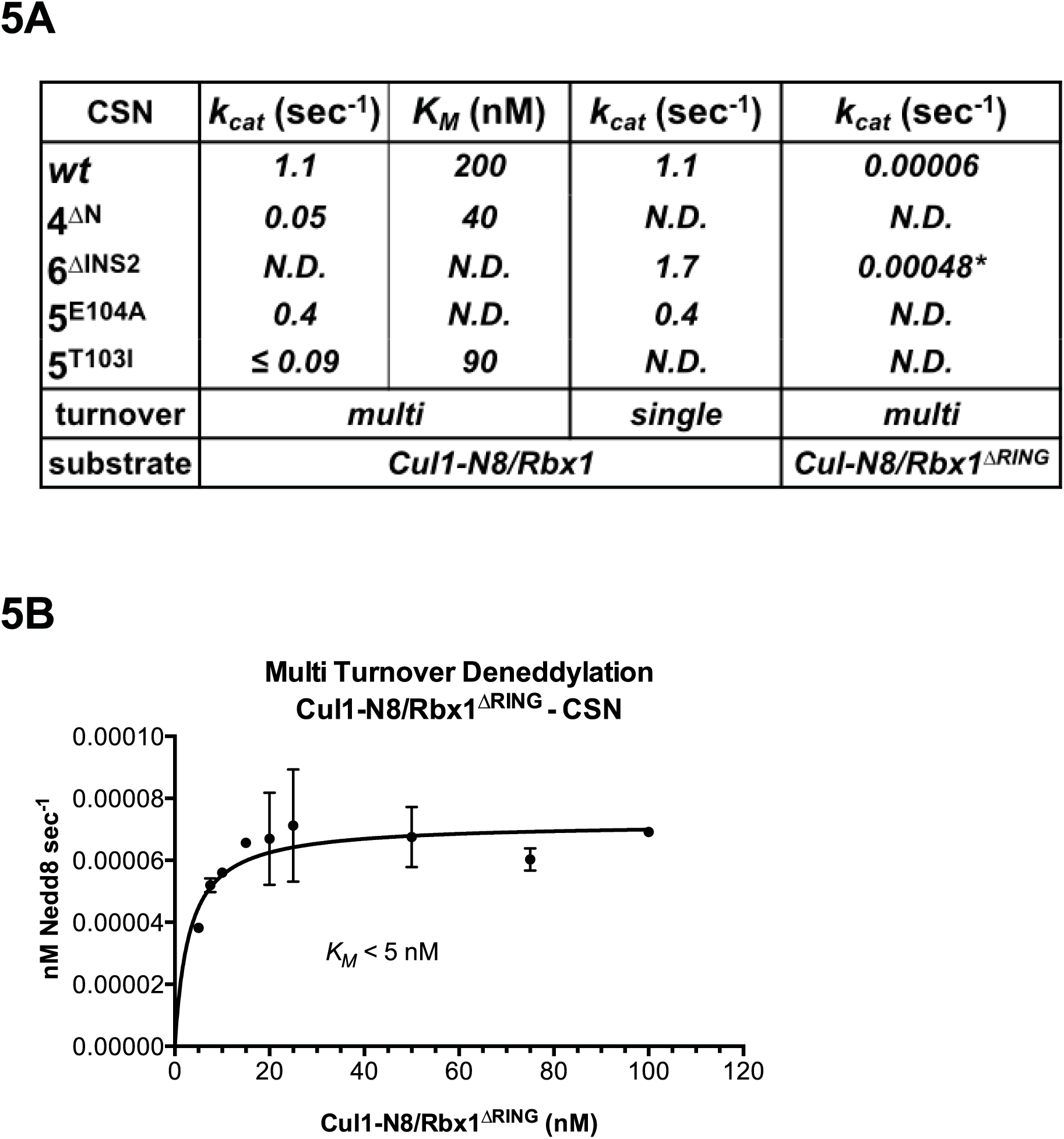
**The N-terminal domains of Csn2 and Csn4 and the RING domain of Rbx1 are important for CSN-mediated deneddylation**. (A) Summary of kinetic parameters for the indicated CSN mutants in multi- or single-turnover deneddylation reactions with Cul1-N8/Rbx1 or Cul1-N8/Rbx1ΔRING substrate. Note that there may be modest discrepancies between these *k_cat_* values and *k_off_* values due to differences in assay configurations as described in Materials and Methods. The ΔRING substrate used here and in panel B contains the sortase sequence at the C-terminus of Cul1 that was used for generation of dansylated Cul1. Control experiments revealed that this tag, with or without dansylation, reduced *k_cat_* by ~4-fold. In addition the wild type control for the ΔRING reaction exhibited *k_cat_* of 2.6 sec^-1^. The rates shown have been correspondingly adjusted to normalize them to other rates reported here. *, This rate is estimated from Figure 5 – figure supplement 5B. (B) Kinetic analysis of deneddylation of Cul1-N8/Rbx1^ΔRING^ by CSN.

A noteworthy feature of the deneddylation reactions carried out with CSN^4Δn^ enzyme or ΔRING substrate is that although *k_cat_* was reduced in both cases, *K_M_* was also reduced (Figure 5A). Whereas these results imply that deletion of the Csn4-NTD or Rbx1-RING improved affinity of the [ES] complex, our direct binding measurements indicated this was not the case. To understand this apparent paradox, it is essential to consider the kinetic behavior of CSN-mediated deneddylation. The formal definition of *K_M_* for a deneddylation reaction ((Equation 1), as stipulated by Briggs and Haldane (Briggs and Haldane, 1925), is: *K_M_* = (*k_off_* + *k_cat_*)/*k_on_*. In the special case of Michaelis-Menten kinetics, which is based on the assumption that *k_off_* is much larger than *k_cat_* the expression simplifies to *k_off_*/*k_on_*, or *K_d_*. However, *k_cat_* for CSN (~1.1 sec^-1^) is actually much faster than *k_off_* measured for dissociation of substrate from the CSN^5E76A,5H138A^ mutant (0.017 sec^-1^). The implication of this is that almost every binding event between CSN and substrate results in catalysis, and *K_M_* (200 nM; Figure 5A and (Emberley et al., 2012) is much larger than *K_d_* (1.6 nM, Figure 3B). But, if *k_cat_* is reduced by mutation, the Briggs-Haldane equation predicts that *K_M_* should approach *K_d_*. Indeed, this is exactly what we see for reactions that exhibit reduced *k_cat_*, including reactions with mutant CSN^4Δn^ enzyme or mutant ΔRING substrate (Figure 5A). In the slowest reaction (cleavage of Cul1/Rbx1^ΔRING^ by CSN) the *K_M_* (5 nM) is in the same range as the *K_d_* with which this substrate bound to CSN^5E76A,H138A^ (12 nM; Figure 3B), and approaches the *K_d_* measured for binding of substrate to CSN^5H138A^ (1.6 nM). This provides strong support for our proposal that the CSN^5H138A^–Cul1^d^/Rbx1 complex is representative of the affinity that develops during normal catalysis.

**Equation 1: Cul1-N8/Rbx1 deneddylation by CSN. The vertical red bar indicates the cleaved bond**.

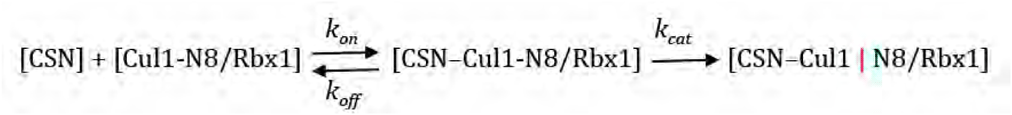

### Functional significance of Csn5 INS1 *in vitro* and in cells

To understand the significance of the E-vict mechansim to CSN function *in vitro* and in cells, we measured the *k_cat_* for CSN^5E104A^ and observed that it is 2.5-fold slower than for CSN (Figure 5A, Figure 5–figure supplements 5C–E). This was unexpected, because it was reported that the Csn5-E104A mutation enhances the catalytic activity of CSN towards an unnatural substrate (Lingaraju et al., 2014). Interestingly, a similar reduced rate was observed in both single- and multi-turnover reactions, indicating that under our specific reactions conditions, the activating conformational changes/chemical step are affected at least as much as product dissociation. This may not be the case in cells, where substrate receptors and other factors may further stabilize product binding.

To test if Csn5-E104 contributes to CSN function *in vivo*, we generated a partial knockout of Csn5 in HEK293T cells using CRISPR/Cas9 (Shalem et al., 2014a). This cell line expressed severely reduced levels of Csn5 and consequently displayed hyper-accumulation of Nedd8-conjugated endogenous Cul1 (Figure 6A), but retained sufficient protein to survive. We introduced either an empty retrovirus or retroviruses coding for Flag-tagged wild type or mutant Csn5 proteins into these cells, and then monitored the Cul1 neddylation status by immunoblotting. In contrast to wild type ^Flag^Csn5, cells expressing ^Flag^Csn5-E104A, H138A or E76A did not regain a normal pattern of Cul1 neddylation (Figure 6A). The same was observed for Cul2, Cul3, Cul4A, and Cul5 (Figure 6– figure supplement 6A). Consistent with reduced CSN activity, as revealed by increased cullin neddylation, Skp2 levels were reduced in cells expressing mutant Csn5 proteins (Figure 6A) (Cope and Deshaies, 2006; Wee et al., 2005). To test whether mutations in the catalytic site of Csn5 resulted in increased affinity for Cul1, we immunoprecipitated wild type and mutant ^Flag^Csn5 proteins and probed for co-precipitation of endogenous Cul1. In addition to the mutants described above, we surveyed a much broader panel of catalytic site substitutions to determine whether the results observed in our *in vitro* experiments were specific to the mutations employed or were a general consequence of disrupting the active site. As shown in Figure 6B, the results were concordant with what was observed *in vitro*. On the one hand, ^Flag^Csn5–H138A retrieved high levels of Cul1-N8. The same was true for ^Flag^Csn5 carrying mutations in other core residues of the JAMM domain (e.g. H140 and D151) (Cope et al., 2002). On the other hand, ^Flag^Csn5-E104A retrieved high levels of unmodified Cul1. We propose that this arises from its ability to bind and deneddylate substrate (albeit at a reduced rate), but then remain tightly bound to the product due to loss of the E-vict mechanism.

**Figure 6.**
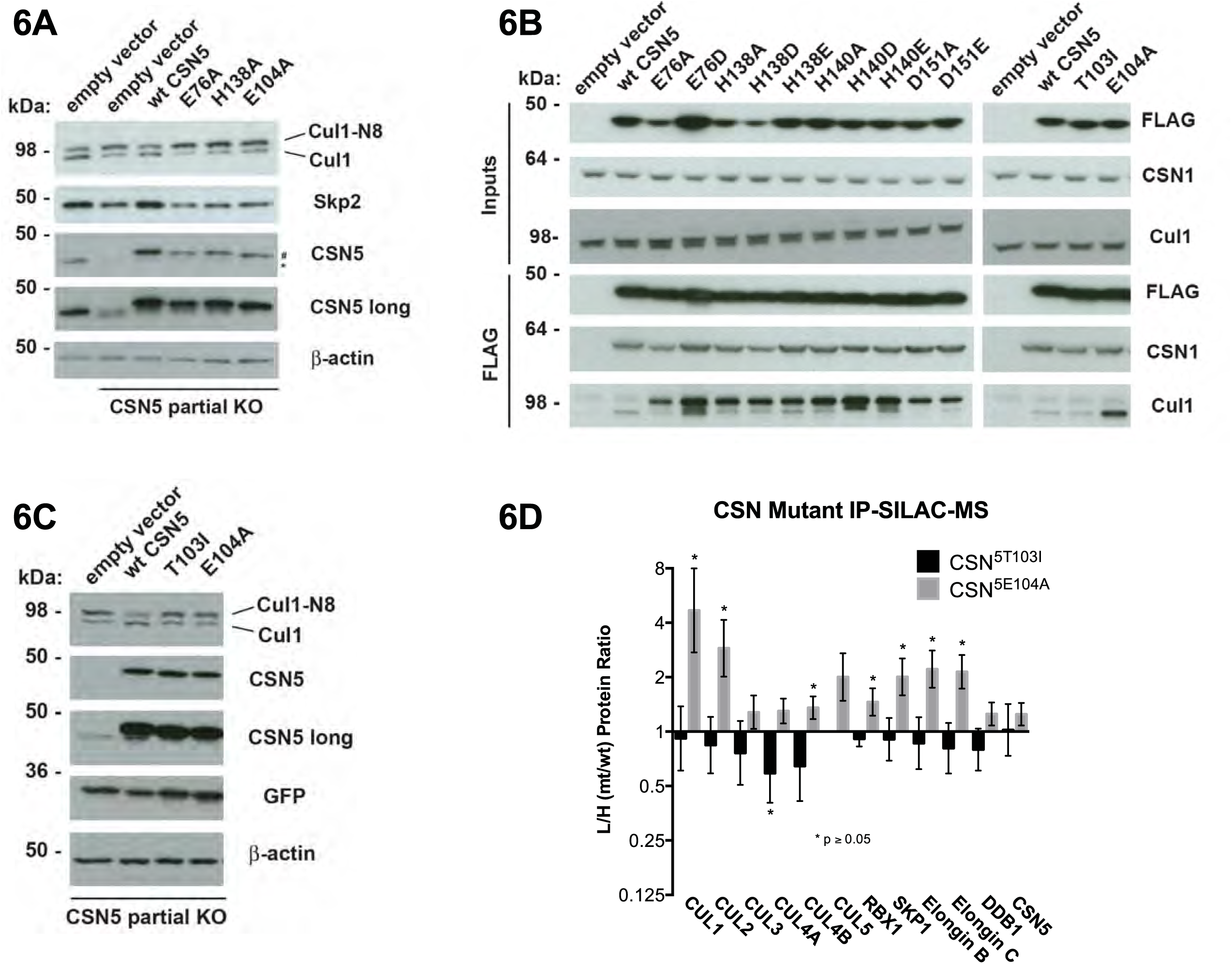
**Functional analysis of Csn5 active site and INS1 mutants in biochemical and cellular assays**. (A) Csn5-E104 is important for CSN function in cells. *CSN5* alleles in HEK293T cells were partially knocked out (KO) by CRISPR/Cas9 to yield a major decrease in Csn5 that was nonetheless compatible with viability. Wild type and the indicated Flag-tagged *CSN5* mutants were reintroduced by transduction of recombinant retroviruses that co-expressed GFP. Lysates of transduced cells were separated by SDS-PAGE and blotted with antibodies to the indicated proteins. CSN5 long refers to a long exposure of the Csn5 blot, captured to reveal residual Csn5 in the knock-out cells. # refers to transduced ^Flag^Csn5 and * refers to endogenous Csn5. (B) Any mutation of a core JAMM domain residue in Csn5 results in enhanced binding to Cul1. Same as (A), except additional Csn5 mutants were tested and the cell lysates were immunoprecipitated with anti-Flag and the immunoprecipitates were blotted for the indicated proteins. (C) Csn5-T103 is important for CSN function in cells. Same as (A) except that the Csn5-T103I mutant was analyzed in parallel with Csn5-E104A and wild type. (D) SILAC mass spectrometry of endogenous proteins bound to ^Flag^Csn5-E104A or ^Flag^Csn5-T103I, relative to wild type ^Flag^Csn5. Cells expressing mutant and wild type ^Flag^Csn5 proteins were grown in light and heavy medium, respectively. L:H ratios >1 indicate higher recovery of the listed protein from cells expressing mutant ^Flag^Csn5, whereas ratios <1 indicate higher recovery from cells expressing wild type ^Flag^Csn5. Gray bars: ^Flag^Csn5-E104A; black bars: ^Flag^Csn5-T103I. Error bars represent the 95% confidence interval as calculated by limma (Smyth, 2005). Each protein was quantified in at least two of the four biological replicates and error bars represent standard deviations. Ratios indicated by * differed significantly from 1.0 (p<0.05). For CSN, only Csn5 is shown; the remainder is shown in Figure 6–figure supplement 6B.

The unexpected reduction in activity observed for Csn5-E104A both *in vitro* and in cells (Figures 5A, 6A) suggested that the adjacent residue, T103, may also be important for deneddylation. A T103I mutation in *Drosophila melanogaster* impedes proper interaction of photoreceptor neurons with lamina glial cells in the developing brain. If this mutant also causes a loss of CSN deneddylase activity, it would explain the recessive nature of this mutation in flies (Suh et al., 2002). Indeed, like Csn5-E104A, Csn5-T103I did not restore a normal Cul1 neddylation pattern when expressed in Csn5-depleted cells (Figure 6C) and exhibited low deneddylase activity *in vitro* (Figures 5A, Figure 5–figure supplement 5F). In contrast to CSN^5E104A^, however, CSN5^T103I^ bound Cul1^d^/Rbx1 product with low affinity, both *in vitro* (Figure 3B, Figure 3–figure supplement 3S) and in cells (Figure 6B). Therefore, although CSN^5E104A^ and CSN^5T103I^ both have diminished catalytic activity, their defects appear to have distinct molecular bases. To further explore the divergent effects of Csn5-E104A and Csn5-T103I mutations on Cul1 binding, HEK293T stably expressing wild type or mutant versions of ^Flag^Csn5 were grown in ‘heavy’ SILAC medium (wild type), or ‘light’ SILAC medium (mutants). Each mutant lysate was individually mixed with wild type lysate, and then subjected to immunoprecipitation and SILAC mass spectrometry. Whereas all CSN subunits exhibited light:heavy ratios of ~1 (Figure 6–figure supplement 6B), the ^Flag^Csn5-E104A pull-down showed elevated levels of all cullins compared to wild type, whereas ^Flag^Csn5-T103I pulled down cullins at levels equal to or less than wild type ^Flag^Csn5 (Figure 6D).

## Discussion

### ‘Induced fit’ underlies interaction of substrate with CSN and triggers enzyme activation

Figure 7 displays a model that incorporates published data and data presented in this manuscript. Panels A-C shows a schematic view of the structural transitions that occur upon substrate binding, and collectively contribute to efficient catalysis, whereas panel D provides the rate constants for the deneddylation cycle. We tentatively propose the following sequence of events. Free CSN exists in an inactive state in which E104 of Csn5-INS1 forms a fourth ligand to zinc (Figure 7A) (Lingaraju et al., 2014). In this state the NTDs of both Csn2 and Csn4 are in “open” conformations relative to the cullin substrate, and the MPN domains of Csn5/Csn6 are in a distal position relative to it. Substrate binds this state rapidly (Figure 7B), likely driven by electrostatic interactions between Cul1 and Csn2-NTD. This would account for the similar, extremely fast *k_on_* values that we measured for different combinations of Cul1^d^/Rbx1 and CSN. Binding of CSN to neddylated substrate results in a series of conformational changes in both complexes (Figure 7B). These include (i) the translocation of the N-terminal helical repeats of Csn2 towards the CTD of Cul1, (ii) the movement of the NTD of Csn4 towards the RING domain of Rbx1 and the WHA domain of Cul1, (iii) the translocation of the MPN domains of Csn5–Csn6 towards the neddylated WHB domain of Cul1, (iv) movement of the WHB domain towards Csn5, (v) the opening of the interface between Nedd8 and the WHB domain, and (vi) the formation of a new interface between Csn5 and Nedd8 probably involving the hydrophobic patch of Nedd8 and neighboring residues, as well as a tenuous interface between the WHB and the Rbx1 RING domain. Furthermore, although not structurally resolved in the present study, movements of Csn5-E76 and E104 towards and away from the zinc atom (vii), respectively, probably similar to the conformation reported in (Echalier et al., 2013), must occur to enable catalysis. Finally, a series of other unresolved movements are likely to be germane including (viii) positioning of the extended C-terminus of Nedd8, and the corresponding portion of the WHB domain for catalysis as well as contacts between the INS1 and INS2 domains of Csn5 and the WHB domain of Cul1.

**Figure 7.**
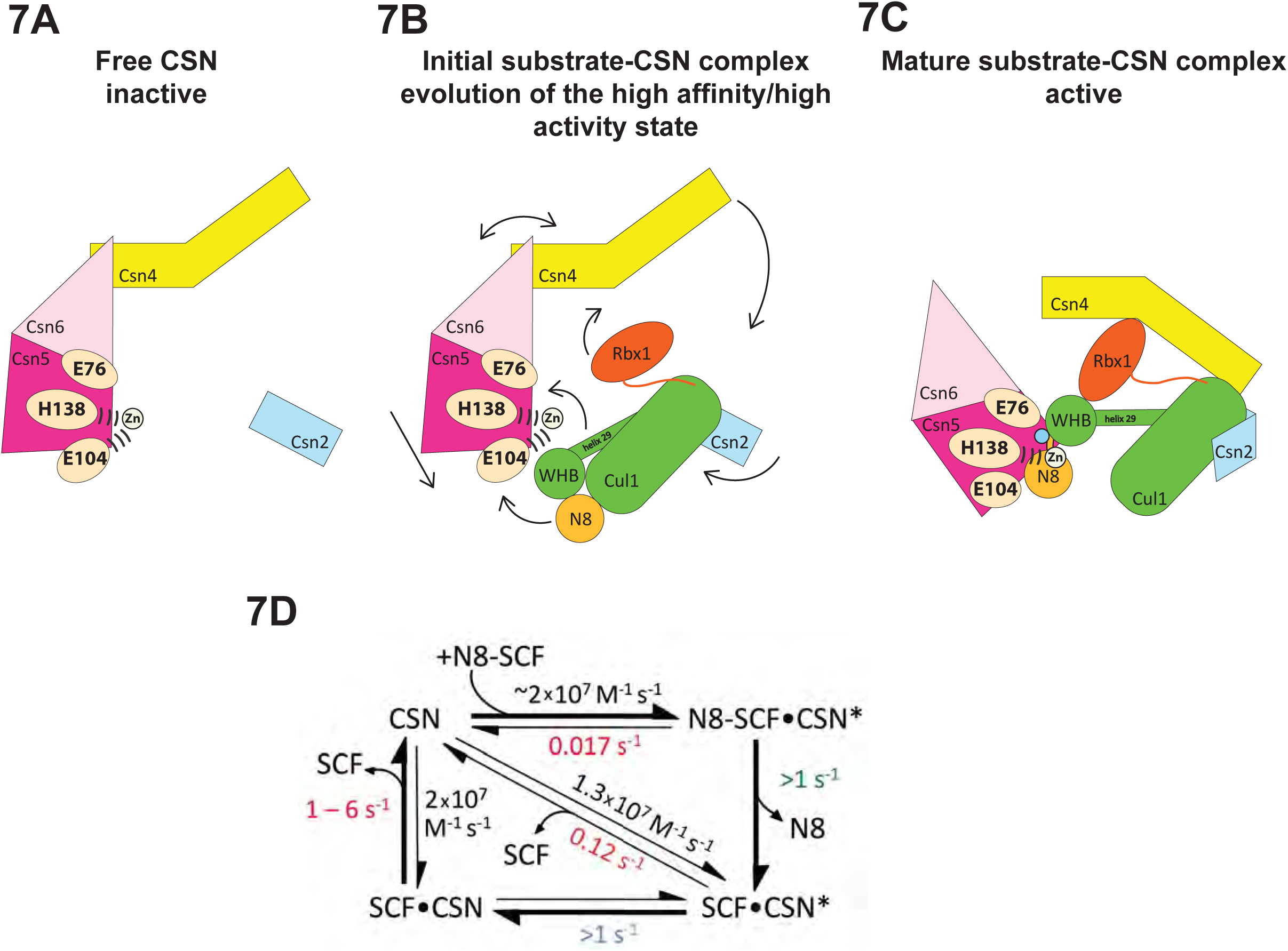
**Structural and kinetic models for CSN activation and the CSN enzyme cycle**. (A-C) Proposed conformational changes that precede substrate cleavage. (D) Kinetic model for the deneddylation cycle. Substrate cleavage is indicated by the slash between N8 and SCF. The asterisk denotes the activated form of CSN. Numbers in red, black, green, and blue represent *k_off_* (sec^-1^), *k_on_* (M-1 sec^-1^), *k_cat_* (sec^-1^), and conformational change (sec^-1^) rates, respectively. For rates >1, the actual rate has not been measured but it is inferred to be >1 sec^-1^ because the overall rate for multiturnover catalysis is at least 1.1 sec^-1^ and thus all sub-steps must be at least this fast. The *k_off_* of SCF from CSN varied depending upon whether the rate was measured directly or inferred from *K_d_* and *k_on_* (see Fig. 3C). The arrow connecting CSN and N8-SCF•CSN* combines two separate steps: binding of N8-SCF to CSN, and activation of CSN to CSN*.

To probe the significance of the conformational changes summarized above, we generated and analyzed mutant enzymes. Deletion of Csn2-NTD virtually eliminated substrate binding (Figure 3–figure supplement 3K), suggesting that movement of this domain (motion i) enables a high affinity interaction between CSN and neddylated CRLs. Meanwhile a mutant lacking Csn4-NTD, CSN^4ΔN,5H138A^, bound Cul1-N8/Rbx1 ~10-fold less tightly than CSN^5H138A^, albeit still with a relatively high affinity (20 nM, Figure 3B). A similar effect on binding affinity was seen with a substrate lacking the RING domain of Rbx1 (Figure 3B). Even though the RING and Csn4-NTD domains are adjacent in the enzyme-substrate [ES] complex (Figure 1), double mutant analysis suggests that they make substantially independent contributions to binding energy (Figure 4B). Interestingly, enzyme assays revealed a much greater effect of deleting the RING than deleting the Csn4-NTD, suggesting that the RING domain makes a profound contribution to catalysis in a manner that does not depend on its proximity to Csn4-NTD. We do not know the extent to which the reduced catalytic rates for these mutants arise from defects in enzyme activation versus substrate positioning, but we note that cleavage of ΔRING substrate was accelerated by ~8-fold upon deletion of Csn6-INS2, suggesting that at least a small part of the problem with this substrate is that it failed to properly trigger activating conformational changes in CSN.

In addition to the movements of individual domains, formation of the [ES] complex is accompanied by wholesale translocation of the Csn5 and Csn6 subunits. We suggest that this motion contributes primarily a *k_cat_* effect, because deletion of Csn6-INS2, which is proposed to facilitate this motion, enhanced *k_cat_* but had no noteworthy impact on binding to substrate (Figure 3B), and removal of Csn5 from the complex did not substantially affect CSN assembly with substrate (Enchev et al., 2012). We cannot conclude much about the other motions enumerated above, but we note that mutations that are predicted to reside near the interface of the Csn5-INS2 and Cul1-WHB domains cause significant reductions in substrate deneddylation (Figure 1–figure supplements 1N–O). In addition, reorientation of Nedd8 away from Cul1-WHB and towards Csn5 as predicted here is consistent with the prior observation that the hydrophobic patch of Nedd8 recruits UBXD7 to neddylated CRLs (den Besten et al., 2012). Presumably, the conformational changes that occur during the activation process are connected in some manner. Interestingly, CSN^5E104A^ and CSN^6Δins2^ both cleave ubiquitin-rhodamine at 0.04 sec^-1^ (which is ~6-fold faster than wild type CSN), but CSN^5E104A,6ΔINS2^ is yet 5-fold faster (0.2 sec^-1^) than either single mutant (Lingaraju et al., 2014). The activities of the single and double mutants imply that the Csn6-ΔINS2 mutation must destabilize binding of Csn5-E104 to the catalytic zinc, but only in a small fraction (≤20%) of complexes. Meanwhile, movements at the Csn4/6 interface must do more to the active site than simply disrupt the interaction of Csn5-E104 with the catalytic zinc, implying the existence of at least two inputs to CSN activation. Resolving how binding of substrate is connected to enzyme activation awaits high-resolution structural analyses of the enzyme and substrate in various states.

### A kinetic model for the CSN enzyme cycle reveals an essential role for the E-vict mechanism in sustaining rapid catalysis

Upon formation of an [ES] complex, the conformational changes that occur in both CSN and substrate culminate in cleavage of the isopeptide bond that links Nedd8 to cullin. Although we don’t know the microscopic rate constants for the various conformational changes and bond cleavage, all evidence points to the former being slower than the latter, which can occur with *k* ≥ 6.3 sec^-1^, based on the *k_cat_* for cleavage of N8-CRL4A^ddb2^ by CSN^6Δins2^ (Lingaraju et al., 2014). The actual cleavage may be even faster because this measurement was made under multi-turnover conditions, in which case product dissociation may have been rate limiting. Regardless, the sum total rate of the activating conformational motions plus isopeptide bond cleavage reported here (~1 sec^-1^) is considerably faster than substrate dissociation from CSN^5H138A^ (~0.017 sec^-1^), indicating that CSN conforms to Briggs-Haldane kinetics and essentially every [ES] complex that forms proceeds to cleavage, the physiological implications of which are considered in the next section.

Cleavage of the isopeptide bond initiates a series of events leading to product release. Removal of Nedd8 increases dissociation of Cul1/Rbx1 by ~7-10 fold. We propose that dissociation of the cleaved Nedd8 also removes an impediment to Csn5-INS1, which can now bind the catalytic site zinc via E104 to return CSN to its apo state. This engagement, which we refer to as the ‘E-vict’ mechanism, is a critical step in what is likely to be a series of conformational rearrangements that include repositioning of Csn5-E76. Collectively, these movements reduce the affinity of CSN for product and accelerate its rate of dissociation by an additional order of magnitude. The removal of Nedd8 and E-vict together bring about an ~100-fold loss in affinity of Cul1/Rbx1 for CSN. The slow dissociation of product from CSN mutants that were unable to undergo E-vict (0.12-0.16 sec^-1^; Figure 3C) suggests that this mechanism is important for maintaining physiological rates of CRL deneddylation. This is further supported by the observation that Csn5-E104A, but not wild type Csn5, co-precipitates substantial amounts of deneddylated Cul1 from cells (Fig. 6B). Slow clearance of product could explain, in part, the failure of this mutant to complement a Csn5 deficiency (Fig. 6A). The E-vict mechanism presents an elegant solution to a fundamental challenge facing enzymes: how to achieve high specificity without compromising rapid turnover.

We note that the product *k_off_* for Cul1^d^/Rbx1 (1.1 sec^-1^) is similar to the *k_cat_* we measured for both single- and multi-turnover reactions. This suggests that depending on the exact structure of the neddylated CRL substrate, the rate-limiting step may vary from one deneddylation reaction to another. Regardless, our biochemical and cell-based data suggest that if the E-vict mechanism did not exist, product dissociation would become the Achilles heel of deneddylation reactions.

### CSN in its cellular milieu

The kinetic parameters reported here coupled with quantitative measurements of protein concentrations by selected reaction monitoring mass spectrometry ((Bennett et al., 2010) and J.R. and R.J.D., unpublished data) allow a preliminary estimate of the steady-state distribution of CSN in cells. The total cullin concentration in the 293T cell line used in this work is ~2200 nM. Meanwhile, the CSN concentration is ~450 nM. Although the total amount of Nedd8-conjugated cullins was not measured, immunoblot data suggest that ~1000 nM is a reasonable estimate. The *K_d_* reported here for the [ES] complex (~2 nM), thus predicts potentially near-complete saturation of the cellular CSN pool with neddylated cullins. This implies that formation of new [ES] complexes is limited by the slowest step in the catalytic cycle, i.e. either the conformational rearrangements or product dissociation. *In vitro*, CSN follows Briggs-Haldane kinetics and cleaves Nedd8 off nearly every neddylated CRL that it binds. Because CSN is not in equilibrium with its substrates in our simplified *in vitro* system, it cannot rely on differences in substrate *K_d_* to achieve specificity. Thus, differences in *k_off_* on the order of ≤10-fold, which might occur with different cullins or substrate adaptors, would be predicted to have minimal effects on catalytic efficiency provided that *k_on_* remains roughly the same, as was observed for different configurations of substrate and product in this study. Importantly, this parameter can potentially be profoundly altered by ubiquitylation substrates, E2 enzyme, or other *in vivo* binding partners of Nedd8-conjugated CRLs, which compete with CSN (Emberley et al., 2012; Enchev et al., 2012; Fischer et al., 2011) and thus should reduce its apparent *k_on_*. It is also conceivable that binding partners might alter the partitioning of the CSN–N8-CRL complex either by increasing *k_off_* and/or reducing *k_cat_*, such that N8-CRL bound to an ubiquitylation substrate dissociates prior to completion of the conformational rearrangements that culminate in its deneddylation.

Based on measurements reported here, it is likely that CSN complexes in cells are constantly undergoing catalysis, dissociating rapidly from product, and rebinding other CRLs on the time-scale of a few seconds or less. Consistent with this picture, addition of a Nedd8 conjugation inhibitor to cells leads to nearly complete disappearance of neddylated cullins within 5 minutes, and this does not account for the time it takes the drug to equilibrate across the membrane and deplete the cellular pool of Nedd8~Ubc12 thioesters (Soucy et al., 2009). The dynamic properties of CSN measured here reveal a CRL network of extreme plasticity that can be reconfigured in minutes to respond to changing regulatory inputs. Although quantitative studies of CRL network dynamics remain in their infancy, it is evident that the tools are at hand to begin to understand how these remarkable enzymes function and are regulated within cells.

## Materials and Methods

### Cloning

All eight wild type CSN subunits were cloned into a single pFBDM baculovirus transfer MultiBac vector (Berger et al., 2004). His_6_-Csn5 was inserted into the first multiple cloning site (MCS1) of a pFBDM vector using NheI/XmaI and Csn1 was put into MCS2 of the same vector with BssHII/NotI. Similarly, Csn2 was inserted into a second pFBDM vector using BssHII/NotI and StrepII^2x^-Csn3, containing an N-terminal PreScission-cleavable StrepII^2x^-tag, using NheI/XmaI. From this plasmid the Csn2/StrepII^2x^-Csn3 gene cassette was excised out with AvrII/PmeI and inserted into pFBDM^Csn1/His6Csn5^, whose multiplication module had been linearized with BstZ17I and SpeI, yielding pFBDM^Csn1/His6-Csn5/Csn2/StrepII2x-Csn3^. A pFBDM^Csn4/Csn7b^ vector was generated using BssHII/NotI to insert Csn4 and NheI/XmaI for Csn7b, and the resultant gene cassette was inserted into linearized pFBDM^Csn1/His6-Csn5/Csn2/StrepII2x-Csn3^, resulting in pFBDM^Csn1/His6-Csn5/Csn2/StrepII2x-Csn3/Csn4/Csn7b^. Finally, a pFBDM^Csn6/Csn8^ vector was generated using BssHII/NotI for Csn6 and NheI/XmaI for Csn8 insertion. Once again the resultant gene cassette was inserted into linearized pFBDM^Csn1/His6-Csn5/Csn2/StrepII2x-Csn3/Csn4/Csn7b^, yielding the full wild type CSN vector pFBDM^Csn1/His6-Csn5/Csn2/StrepII2x-Csn3/Csn4/Csn7b/Csn6/Csn8^. A similar cloning strategy was applied for the generation of CSN^5E76A^, CSN^5E76A,H138A^, CSN^5E212R,D213R^ and CSN^4ΔN1-297^, except that site-directed mutageneses were performed on pFBDM^Csn1/His6Csn5^ and pFBDM^Csn4/Csn7b^ respectively. CSN^5E104A^ and CSN^5T103I^ were generated with the same general approach, except that that site-directed mutagenesis and sequence validation were performed on a pCRIITOPO plasmid (Invitrogen) containing StrepII^2x^-Csn5. Those mutants were then ligated into a MCS1 linearized pFBDM^Csn1^ plasmid. For the production of CSN^6Δins2^ we used co-expression from two separate viruses. To this end we applied site-directed mutagenesis on the pFBDM^Csn6/Csn8^ vector to delete amino acids 174-179 in Csn6, generating pFBDM^Csn6ΔIns2/Csn8^. The gene cassette of the latter was excised out using AvrII/PmeI and inserted into BstZ17I/SpeI linearized pFBDM^Csn4/Csn7b^, yielding pFBDM^Csn4/Csn7b/Csn6ΔIns2/Csn8^. The resultant bacmid was used together with a bacmid generated from pFBDM^Csn1/His6-Csn5/Csn2/StrepII2x-Csn3^ in order to generate two baculoviruses, which were used for co-infection to generate CSN^6Δins2^. An analogous strategy was applied to generate CSN^4Δn/6Δins2^, CSN^5H138A/6ΔINS2^ and CSN^5H138A/4ΔN^.

The TEV site in Rbx1 as well as mutations in the WHB domain of Cul1 were obtained by site-directed mutagenesis on the pFBDM-Cul1/Rbx1 vector described in (Enchev et al., 2010), which further contained a C-terminal sortase tag described in the next section. Cloning of Cul3/Rbx1 used in the crosslinking/mass spectrometry experiments, Nedd8-pro-peptide-StrepII^2x^ and StrepII^2x^-Den1 are described in (Orthwein, 2015). Recombinant bacmid and virus generation as well as protein expression proceeded as described in (Enchev et al., 2012). All genes were validated by sequencing as wild type or mutant.

### Protein Purification and modifications

CSN and its mutant forms were purified as described in Enchev et al. (2012). Nedd8-activating and conjugating enzymes were purified as described in Emberley et al. (2012) and Enchev et al. (2012). Fluorescently-labeled Cul1 substrates were conjugated with untagged Nedd8. Cul1-sortase was designed with GGGGSLPETGGHHHHHH inserted after the final amino acid of Cul1 into the pGEX vector described in Emberley et al. (2012). All sortase reactions were done at 30 °C overnight with 30 μM Cul1/Rbx1, 50 μM Sortase and 250 μM GGGGK-dansyl in 50 mM Tris pH 7.6, 150 mM NaCl and 10 mM CaCl_2_ and purified by size exclusion chromatography to yield Cul1^d^/Rbx1. Cul1^d^/Rbx1 was neddylated and purified as in Emberley et al. (2012) to yield Cul1^d^-N8/Rbx1. Cand1 and Sortase were purified as described in Pierce et al. (2013). Production of Cul1/Rbx1 and Cul3/Rbx1 baculovirus constructs used for electron microscopy and crosslinking mass spectrometry, bacterial split-and-co-express Cul1/Rbx1^ΔRING^, Nedd8 with native N- and C-termini, used for electron microscopy and crosslinking mass spectrometry and for the experiments involving Cul1/Rbx1^TEV^, Den1 as well as the respective preparative neddylation were performed as described in Enchev et al. (2012) and (Orthwein et al., 2015). Den1 was used in 1:50 ratio for 10 min at 25 °C to remove poly-neddylation. Cul1/Rbx1 complexes with mutations in the WHB domain of Cul1 (Figure 1–figure supplement 1N, O) and Cul1/Rbx1^TEVΔRING^ were purified from High Five insect cells as described in Enchev et al. (2010). Dansylation of Cul1/Rbx1 variants expressed in insect cells was performed for 8 to 12 h at 30 °C while spinning at 5000 *g*, and purified by passing the dansylation reaction through a 5 ml HisTrap FF column (GE Healthcare) in 50 mM Tris-HCl, pH 7.6, 400 mM NaCl, 20 mM imidazole. The Cul1^d^/Rbx1-containing flow through was concentrated, neddylated (if required), and further purified over a Superdex 200 size exclusion column (GE Healthcare) equilibrated with 15 mM HEPES, pH 7.6, 150 mM NaCl, 2 mM DTT, 2 % (v/v) glycerol. Neddylation of Cul1/Rbx1^TEVΔRING^ was performed at 25 °C for 12-14 h in 50 mM Tris-HCl, pH 7.6, 100 mM NaCl, 2.5 mM MgCl_2_, 150 μM ATP, spinning at 2000 *g*, and was followed by 30 min incubation with 1:50 (w/w) Den1 to remove poly-neddylation. The reaction was purified over a Strep-Tactin Superflow Cartridge (QIAGEN), and eluted in 15 mM HEPES, pH 7.6, 250 mM NaCl, 2 mM DTT, 2 % (v/v) glycerol, 2.5 mM *d*-desthiobiotin. RING cleavage was performed for 12-14 hours at 25 °C, spinning at 2000 *g*, in the presence of 100 mM EDTA, pH 8 and 1:1 (w/w) TEV. Dansylation proceeded as described above.

### Deneddylation Assays

All deneddylation assays were performed in a buffer containing 25 mM Tris-HCl, pH 7.5, 100 mM NaCl, 25 mM trehalose, 1 mM DTT, 1 % (v/v) glycerol, 0.01 % (v/v) Triton X-100 and 0.1 mg/ml ovalbumin or BSA. Radioactive deneddylation reactions with bacterially expressed substrates were done as described (Emberley et al., 2012). Radioactive deneddylation reactions with substrates expressed in insect cells were performed at 24 °C with 0.5 nM CSN (Figure 2–figure supplement 2C) or 2 nM CSN (Figure 5B). All remaining radioactive deneddylation reactions were performed with bacterially expressed Cul1-N8/Rbx1 substrates (50 nM) and 2 nM CSN unless otherwise noted. Single-turnover reactions were done with 25 nM Cul1 substrates and 1 μM CSN on a Kintek RQF-3 Rapid Quench Flow at 24 °C. Single-turnover data were fit to one phase decay function: Y = (Y0 - EP)*exp(-*k_cat_*\*X) + EP (where EP corresponds to reaction end point value), to determine the *k_cat_*. Deneddylation assays in Figure 1–figure supplement 1N, O were performed with 800 nM substrate and 20 nM enzyme and visualized by Coomassie stain. Depending on the exact protein preparations used and the laboratory, we observed rates for the wild type reaction ranging from 1.1-2.6 sec^-1^.

### Fluorescence Assays

All assays were performed in a buffer containing 30 mM Tris pH 7.6, 100 mM NaCl, 0.25 mg/ml ovalbumin or BSA and 0.5 mM DTT with 30 nM dansyl-labeled Cul1/Rbx1 and titrated concentrations of CSN. The mixtures were allowed to reach equilibrium by incubation at room temperature for ~ 10 minutes prior to measurements. Equilibrium binding assays using Cul1/Rbx1 variants expressed in insect cells (Figure 2, Figure 2–figure supplement 2E, Figure 3–figure supplement 3N, 3R, Figure 4B) were read at 530 nm on a CLARIOstar plate reader (BMG Labtech) in 384-well plates (Corning, low flange, black, flat bottom), 90 ul per well, while binding assays using bacterially expressed Cul1/Rbx1 variants were performed on a Fluorolog-3 (Jobin Yvon) (all other binding data figures). Binding assay with Cul1^d^-N8/Rbx1 (substrate) and CSN^5E76A^ were allowed to equilibrate for only 45 seconds, because although this mutant exhibited an ~300-fold decrease in activity (data not shown) the residual activity was high enough to cause substantial deneddylation in a 10 minute incubation. It should be noted that several of the *K_d_* values reported for CSN binding to Cul1^d^-N8/Rbx1 or Cul1^d^/Rbx1 are below the concentration of the dansylated ligand (30 nM). While this is generally not the preferred approach, we found that 30 nM was the lowest concentration that consistently yielded highly reproducible results. The estimated *K_d_* is very sensitive to the density of data points at the inflection point of the curve, and thus these estimates can be more prone to error. Nevertheless, different investigators in Zurich and Pasadena have consistently obtained an estimate of 1.6-5 nM for binding of CSN^5H138A^ to Cul1^d^-N8/Rbx1 and of 9-13 nM for binding to CSN^5H138A^ to Cul1^d^/Rbx1, using different protein preparations. To estimate *K_d_*, all data points were fitted to a quadratic equation, Y = Y0+(Ymax-Y0)*(*K_d_*+A+X-sqrt((*K_d_*+A+X)^2-4A*X))/2*A where A equals concentration of labeled protein, using Prism (Graph Pad). On-rate and off-rate measurements were performed on a Kintek Stopped-flow SF-2004 by exciting at 340 nM and collecting emissions through a 520 +/- 20 nm filter. For off-rate measurements, the concentrations of proteins used in each reaction are provided in the legend of Figure 3–figure supplements 3F–I. Off rate data were fit to one phase decay function: Y=(Y0 - EP)*exp(-*k_off_*\*X) + EP (where EP corresponds to reaction end point value).Whereas *K_d_*, on-rate, and off-rate measurements with different configurations of Cul1 or different CSN mutants are directly comparable, off-rate measurements are not directly comparable to *k_cat_* measurements and may differ from expectation by a few fold because different buffers were used, the Cul1/Rbx1 preparations were from different sources (bacterial for *k_cat_*, baculoviral for *k_off_*), and the Cul1/Rbx1 preparations carried different labels (dansylated Cul1 for *k_off_*, [^32^P]-Nedd8 for *k_cat_*.

### Cell Culture and SILAC Mass Spectrometry

Cells were grown in Lonza DMEM containing 10% FBS (Invitrogen). Transient transfections were done with FugeneHD per the manufacturers instructions (Roche). Flag-tagged *CSN5* coding sequences were cloned into a modified MSCV-IRES-GFP vector (containing a pBabe multiple cloning site) via BamHI and EcoRI. Lenti-CRISPR constructs were made as described (Shalem et al., 2014b) using the targeting sequences 5’-CACCGCTCGGCGATGGCGGCGTCC - 3’ and 3’ - AAACGGACGCCGCCATCGCCGAGC - 5’. Lenti- and retroviruses were produced in 293T cells and the supernatant subsequently used for transduction to establish stable cell lines. For Western Blot analysis cells were directly lysed in 2X SDS sample buffer. Lysates were sonicated for 15 seconds at 10 % of maximum amplitude using a Branson Digital Sonifier and boiled for 10 minutes at 100 ºC. SILAC labeling was in Invitrogen DMEM containing 10% FBS and ^13^C_6_^15^N_2_-lysine and ^13^C_6_-arginine from Cambridge Isotope Laboratory. For immunoprecipitations, cells were lysed in Pierce Lysis Buffer containing cOmplete Protease Inhibitor Cocktail (Roche) and lysates were sonicated for 10 seconds at 10 % of maximum amplitude using a Branson Digital Sonifier. After a 5 minute clearing at 18000 x g at 4°C, proteins were immunoprecipitated with M2 Flag agarose beads (Sigma) for 30 minutes and prepared for mass spectrometry as described in Pierce et al. (2013). Samples were analyzed using an EASY-nLC 1000 coupled to an Orbitrap Fusion and analyzed by MaxQuant (v 1.5.0.30).

Digested peptides (250 ng) were loaded onto a 26-cm analytical HPLC column (75 μm ID) packed in-house with ReproSil-Pur C_18_AQ 1.9 μm resin (120 Å pore size, Dr. Maisch, Ammerbuch, Germany). After loading, the peptides were separated with a 120 min gradient at a flow rate of 350 nL/min at 50°C (column heater) using the following gradient: 2-6% solvent B (7.5 min), 6–25% B (82.5 min), 25-40% B (30min), 40-100% B (1min), and 100% B (9 min) where solvent A was 97.8% H_2_O, 2% ACN, and 0.2% formic acid) and solvent B was 19.8% H_2_O, 80% ACN, and 0.2% formic acid. The Orbitrap Fusion was operated in data-dependent acquisition (DDA) mode to automatically switch between a full scan (*m*/*z*=350–1500) in the Orbitrap at 120,000 resolving power and a tandem mass spectrometry scan of Higher energy Collisional Dissociation (HCD) fragmentation detected in ion trap (using TopSpeed). AGC target of the Orbitrap and ion trap was 400,000 and 10,000 respectively.

### SILAC MS data analysis

Thermo RAW files were searched with MaxQuant (v 1.5.3.8) (Cox and Mann, 2008; Cox et al., 2011). Spectra were searched against human UniProt entries (91 647 sequences) and a contaminant database (245 sequences). In addition, spectra were searched against a decoy database (generated by reversing the target sequences) to estimate false discovery rates. Trypsin was specified as the digestion enzyme with up to two missed cleavages allowed. Variable modifications included oxidation of methionine and protein N-terminal acetylation. Carboxyamidomethylation of cysteine was specified as a fixed modification. SILAC was specified as the quantitation method with Arg6 and Lys8 specified as the heavy labeled amino acids. Precursor mass tolerance was less than 4.5 ppm after recalibration and fragment mass tolerance was 0.5 Da. False discovery rates at the peptide and protein levels were less than 1% as estimated by the decoy database search. Ratios were calculated for proteins quantified in at least two of the four biological replicates. 95% confidence intervals and adjusted p-values were calculated using the R package limma (Ritchie et al., 2015)

### Cross-linking coupled to mass spectrometry (XL-MS)

Chemical cross-linking of purified complexes was performed using DSS H_12_/D_12_ (Creative Molecules) as cross-linking agent and as previously described (Birol et al., 2014). Subsequent MS analysis and cross-link assignment and detection were carried out essentially as described (Leitner et al., 2014) on an Orbitrap Elite (Thermo Scientific) using the *xQuest/xProphet* software pipeline.

### Western Blot Analysis

Proteins were separated by SDS-PAGE gel electrophoresis and transferred to a nitrocellulose membrane by wet blot. Primary antibodies used for detection were: anti-CSN5 mouse monoclonal Santa Cruz Biotechnology sc-393725, anti-Cul1 mouse monoclonal Santa Cruz Biotechnology sc-17775, anti-Cul2 rabbit polyclonal Thermo Scientific #51-1800, anti-Cul3 rabbit polyclonal Cell Signaling #2769, anti-Cul4A rabbit polyclonal Cell Signaling #2699, anti-Cul5 rabbit polyclonal Bethyl Laboratories A302-173A, anti-β-actin mouse monoclonal Sigma A5316, anti-GFP mouse monoclonal Clontech #632381.

### Sample preparation for electron microscopy and data collection

CSN^5H138A^-SCF-Nedd8^Skp2/Cks1^samples for cryo-electron microscopy were generated by pre-incubating the purified components as described in Enchev et al (2012) and ran over a Superose 6 increase 3.2/300 column (GE Healthcare) at 4 °C, eluting 50 µl fractions in 15 mM HEPES, pH 7.6, 100 mM NaCl, 0.5 mM DTT. The sample was kept on ice and its homogeneity and mono-dispersity from the peak elution was immediately confirmed by visualization in negative stain. For cryo EM preparation, the sample was diluted to 0.1 mg/ml and 2 µl were applied to Quantifoil grids (R1.2/1.3 Cu 400 mesh), freshly coated with an extra layer of thin carbon and glow-discharged for 2 min at 50 mA and 0.2 mbar vacuum. The grids were manually blotted to produce a thin sample film and plunge-frozen into liquid ethane. Data were collected automatically using EPU software in low dose mode on a Titan Krios transmission electron microscope, equipped with a Falcon II direct electron detector (FEI), and operated at 300 kV, an applied nominal defocus from −2.5 to - 5.0 µm in steps of 0.25 µm, and 80,460-fold magnification, resulting in a pixel size of 1.74 Å on the sample scale. Images were collected as seven separate frames with a total dose of 25 e^-^/Å^2^.

### Electron microscopy data analysis

CTF-estimation and subsequent correction were performed using RELION (Scheres, 2012) and CTFFIND3 (Mindell and Grigorieff, 2003). All micrographs were initially visually inspected and only those with appropriate ice thickness as well as Thon rings in their power spectra showing regularity and extending to 6 Å or beyond were used for subsequent analysis. In order to generate 2D references for automated particle selection, ~ 4,000 single particles were manually picked and subjected to 2D classification in RELION. Six well-defined 2D class averages were selected, low-pass filtered to 35 Å to prevent reference bias, and used as references. Approximately 150,000 single particles were automatically selected and subjected to reference-free 2D and 3D classification, in order to de-select the particles, which resulted in poorly defined or noisy averages. Approximately half of these single particles resulted in a well-defined 3D class average, which resembled the previously published negative stain EM map of the same complex (Enchev et al., 2012). This dataset was subject to 3D auto-refinement in RELION, using a version low-pass filtered to 50 Å as an initial reference. The converged map was further post-processed in RELION, using MTF-correction, FSC-weighting and a soft spherical mask with a 5-pixel fall-off.

### Modeling, docking and visualization

Csn7b was modeled using Csn7a as a template on the Phyre2 server (Kelley et al., 2015) and the modeled coordinates were aligned to Csn7a in PDB ID 4D10 (Lingaraju et al., 2014), effectively generating a CSN atomic model for the Csn7b-containing complex. Model visualization, molecular docking, distance measurements and morph movie generation were performed with UCSF Chimera (Pettersen et al., 2004).

### Accession code

The cryo electron microscopy density map of CSN^Csn5H138A^-SCF-Nedd8^Skp2/Cks1^ is deposited in the Electron Microscopy Data Bank under accession code EMD-3401.

## Acknowledgements

We thank B. Schulman for Skp1/Skp2 protein as well as for expression plasmids and E. Morris for thin carbon for EM grids. We thank A. Bernini and A. Ragheb for technical assistance with protein work, P. Tittmann for technical support at the ScopeM facility, members of the Ban lab for advice and Annie Moradian and Roxana Eggleston-Rangel for mass spectrometry support at PEL. We also are grateful to S.O. Shan, E. Morris and T. Stuwe for advice and A. Smith, S.O. Shan, and D. Barford for comments on the manuscript. R.M. was supported by a Lee-Ramo Life Sciences Fellowship, R.I.E. was supported by an ETH Pioneer, a Marie Curie and an EMBO short-term fellowship, and A.S. by a Marie Curie fellowship. F.S. acknowledges funding from the Wellcome Trust (Grant 095951) and the German Science Foundation Collaborative Research Center (SFB) 969. The Peter laboratory is funded by an ERC advanced grant, the SNF and ETHZ, and the Aebersold laboratory is supported by ETH Zurich, SystemsX.ch and an ERC advanced grant. This work was supported in part by NIH CA164803 to R.J.D. R.J.D. is an Investigator of and was supported by the Howard Hughes Medical Institute. M.J.S and S.H were supported by the Gordon and Betty Moore Foundation, through Grant GBMF775 and the Beckman Institute.

---

**Figure 1–figure supplement 1.**
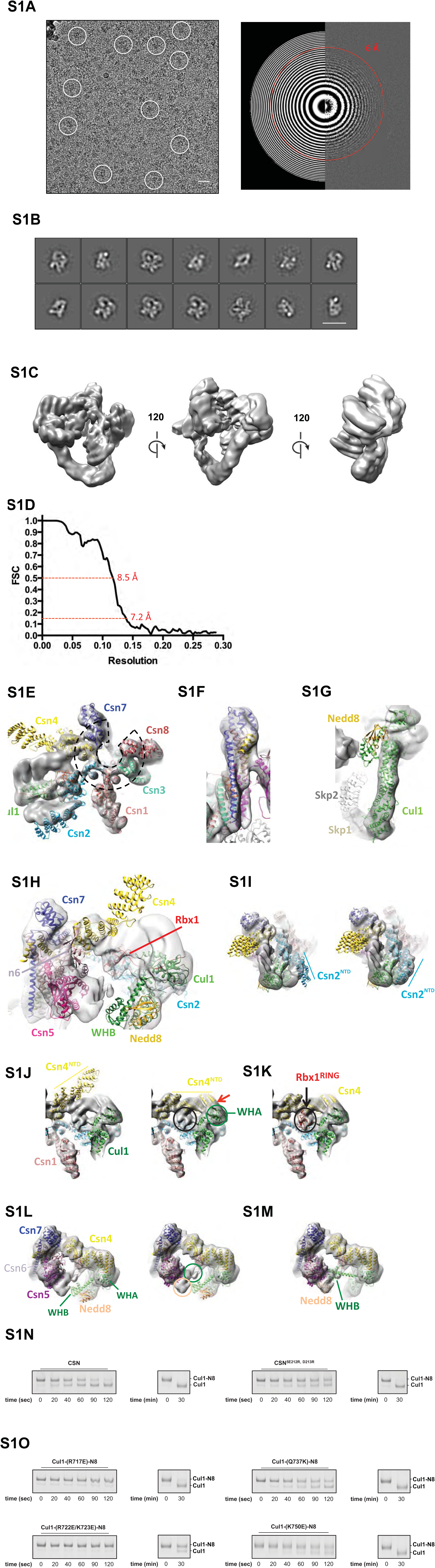
**Cryo-electron microscopy and single particle analysis of a CSN^5H138A^-N8-SCF^Skp2/Cks1^ complex**. (A) A representative cryo-electron micrograph of a CSN^5H138A^-N8-SCF^Skp2/Cks1^ complex with some single molecular views indicated by white circles (left) and a power spectrum indicating Thon rings reaching 6 Å (right). Scale bar is 200 Å. (B) Representative two-dimensional class averages from the curated dataset, used for the subsequent analysis. Scale bar is as in (A). (C) Surface views of the final, post-processed cryo-electron map. (D). Resolution estimate according to the FSC criteria of 0.143 and 0.5. (E) Fit of the PCI-domain containing CSN subunits in the cryo-electron density map. Csn1,3,7, and 8 match the density very well but the N-terminal domains of Csn2 and Csn4 do not, but their winged-helix domains fit well. The horseshoe arrangement of the six winged-helix domains is indicated with a dotted black line. (F) All the C-terminal helices of the CSN subunits match well the electron density map. (G) Fit of SCF in the electron density map. (H) Same view as in Fig 1B but prior to flexible docking of the N-terminal domains of Csn2 and Csn4, the MPN domains of Csn5&6, the WHB domain of Cul1, and Nedd8. (I) Movement of the N-terminus of Csn2 from its crystallographically-determined position (left) into the EM density map (right). (J) Movement of the N-terminus of Csn4 from its crystallographically-determined position (left) into the EM density map (right). The two N-terminal helical repeats of Csn4, red arrow, are in close proximity to the WHA domain of Cul1 (green circle). (K) Localization of the RING domain of Rbx1. The unfilled density that is indicated by a black ellipse in the right-hand panel of S1J accommodates Rbx1 (shown in red). The helices of Csn4 in close proximity to the RING domain of Rbx1 are indicated by a black arrow. (L) Re-localization of Csn5/6. Comparing the left and right panels, Csn5/6 move leftward to occupy unfilled density. The tan and green circles below Csn5/6 indicate densities that are occupied by Nedd8 and the WHB domain, as depicted in (M). (N, O) Deneddylation assays with (N) wild type Cul1-N8/Rbx1 and indicated CSN variants and (O) wild type CSN and mutant Cul1 variants. Note that all Cul1 constructs have an uncleaved C-terminal sortase tag, which is the reason for slower deneddylation of wild type Cul1-N8/Rbx1 by wild type CSN relative to the kinetics reported elsewhere in this work.

**Figure 2–figure supplement 1.**
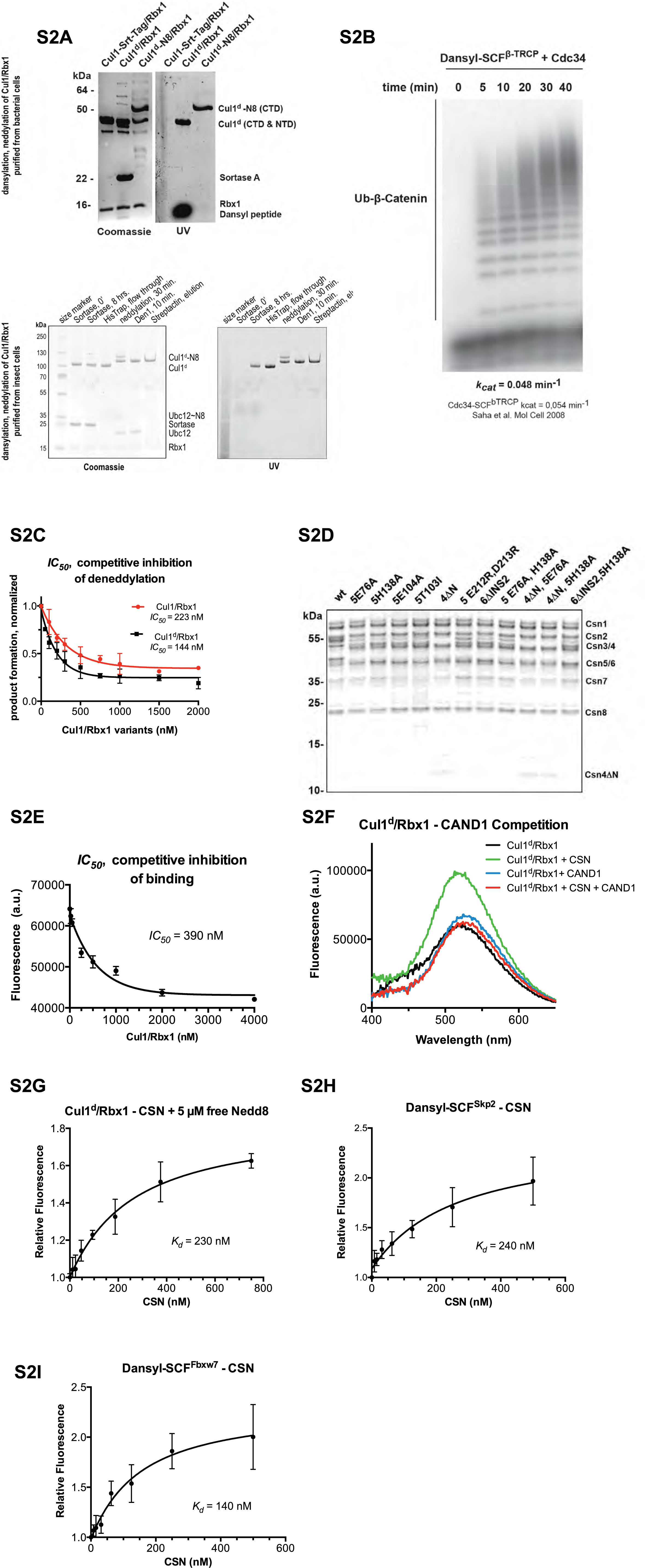
**Supporting data for development and validation of CSN– Cul1^d^/Rbx1 binding assay**. (A) Dansylation of Cul1/Rbx1 constructs. Upper panel: dansylation of bacterially expressed and purified Cul1/Rbx1. Lower panel: dansylation of Cul1/Rbx1 expressed and purified from insect cells. For details, see Materials and Methods. (B) Ubiquitination of 32P-labeled β-catenin substrate peptide by dansylated SCF^β-TrCP^ was monitored as described (Saha and Deshaies, 2008). The *k_cat_* measured here (0.048 min^-1^) compares favorably with that previously determined for wild type unmodified SCF (0.054 min^-1^) (Saha and Deshaies, 2008). (C) *IC_50_* study of the inhibitory effects of unlabeled (red) or dansylated (black) product. Cul1/Rbx1 and Cul1^d^/Rbx1 were separately titrated into a deneddylation reactions containing 50 nM Cul1-[32P]N8/Rbx1 substrate and 0.5 nM CSN, and the resulting reaction rate was measured. (D) CSN preparations used in this study. 600 ng of each sample were fractionated by SDS-PAGE and stained with SYPRO Ruby. (E) *IC_50_* for competitive inhibition of CSN-Cul1^d^/Rbx1 complex formation by unlabeled Cul1/Rbx1 (~ 390 nM) agrees with the *K_d_* measured for binding of Cul1^d^/Rbx1 to CSN (310 nM). (F) Equilibrium binding of 100 nM CSN to 50 nM Cul1^d^/Rbx1 and competition by 500 nM Cand1. The indicated proteins were mixed and allowed to equilibrate prior to determination of dansyl fluorescence. (G-I) Free Nedd8 and F-box box proteins do not appreciably change affinity of Cul1^d^/Rbx1 for CSN. Same as Figure 2C, except that either 5 µM free Nedd8 (G), 100 nM Skp2/Skp1 (H) or 100 nM Fbxw7/Skp1 (I) was included in the binding reaction. All binding and activity measurements reported in this legend were carried out in triplicate and error bars represent standard deviation.

**Figure 3–figure supplement 1.**
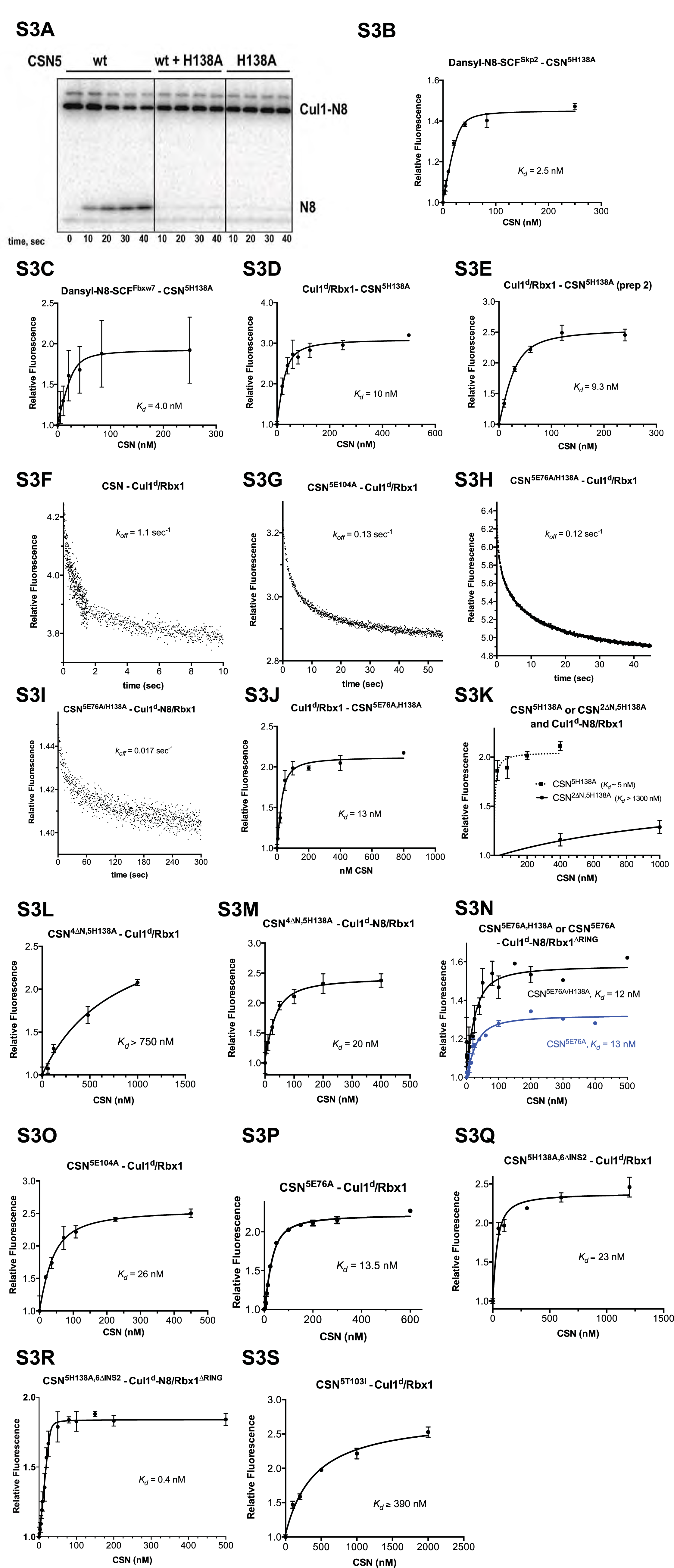
**Supporting experiments and titration curves for binding data in Figures 3B–C**. (A) CSN^5h138A^ is inactive and is a dominant-negative inhibitor of deneddylation. CSN, CSN^5H138A^, and substrate were used at 2 nM, 100 nM, and 75 nM, respectively. For reactions containing with CSN and CSN^5H138A^, mutant enzyme was preincubated with substrate for 30 sec prior to initiating time-course by adding CSN. (B-E): The indicated proteins were mixed and allowed to equilibrate prior to determining the change in dansyl fluorescence. (B) CSN^5H138A^ and dansylated, Nedd8-conjugated SCF^skp2^. (C) CSN^5H138A^ and dansylated, Nedd8-conjugated SCFFbxw7. Note that addition of Fbxw7–Skp1 greatly increased the variability in the measurement for unknown reasons. (D) CSN^5H138A^ (first prep) and Cul1^d^/Rbx1, (E) CSN^5H138A^ (second prep) and Cul1^d^/Rbx1, (F-I): The indicated CSN complexes were preincubated with Cul1^d^/Rbx1 for 10 min, followed by addition of unlabeled Cul1/Rbx1 chase and measurement of the decay in dansyl fluorescence over time. Final protein concentrations are listed for each experiment. (F) CSN (2000 nM), Cul1^d^/Rbx1 (200 nM), and Cul1/Rbx1 (3000 nM), (G) CSN^5E104A^ (600 nM), Cul1^d^/Rbx1 (200 nM), and Cul1/Rbx1 (3000 nM), (H) CSN^5E76A,5H138A^ (400 nM), Cul1^d^/Rbx1 (200 nM), and Cul1/Rbx1 (3000 nM), (I) CSN^5E76A,5H138A^ (200 nM), Cul1^d^-N8/Rbx1 (100 nM), and Cul1/Rbx1 (1500 nM), (J-S): The indicated proteins were mixed and allowed to equilibrate prior to determining the change in dansyl fluorescence. (J) CSN^5E76A,5H138A^ and Cul1^d^/Rbx1, (K) CSN^5H138A^ or CSN2Δn,^5H138A^ and Cul1^d^-N8/Rbx1, (L) CSN^4ΔN,5H138A^ and Cul1^d^/Rbx1, (M) CSN^4ΔN,5H138A^ and Cul1^d^-N8/Rbx1, (N) CSN^5E76A^ or CSN^5E76A,5H138A^ and Cul1^d^-N8/Rbx1^ΔRING^, (O) CSN^5E104A^ and Cul1^d^/Rbx1, (P) CSN^5E76A^ and Cul1^d^/Rbx1, (Q) CSN^5H138A,6ΔINS2^ and Cul1^d^/Rbx1, (R) CSN^5H138A,6ΔINS2^ and Cul1^d^-N8/Rbx1ΔRING, (S) CSN^5T103I^ and Cul1^d^/Rbx1. All measurements in panels B-E and J-S were carried out in triplicate and error bars represent standard deviation. The measurement in panel P was performed in duplicates but the experiment was repeated on three independent occasions, obtaining similar results. Several of these results were independently confirmed in Zurich and Pasadena including panels J, M, O, P, and Q.

**Figure 4–figure supplement 1.**
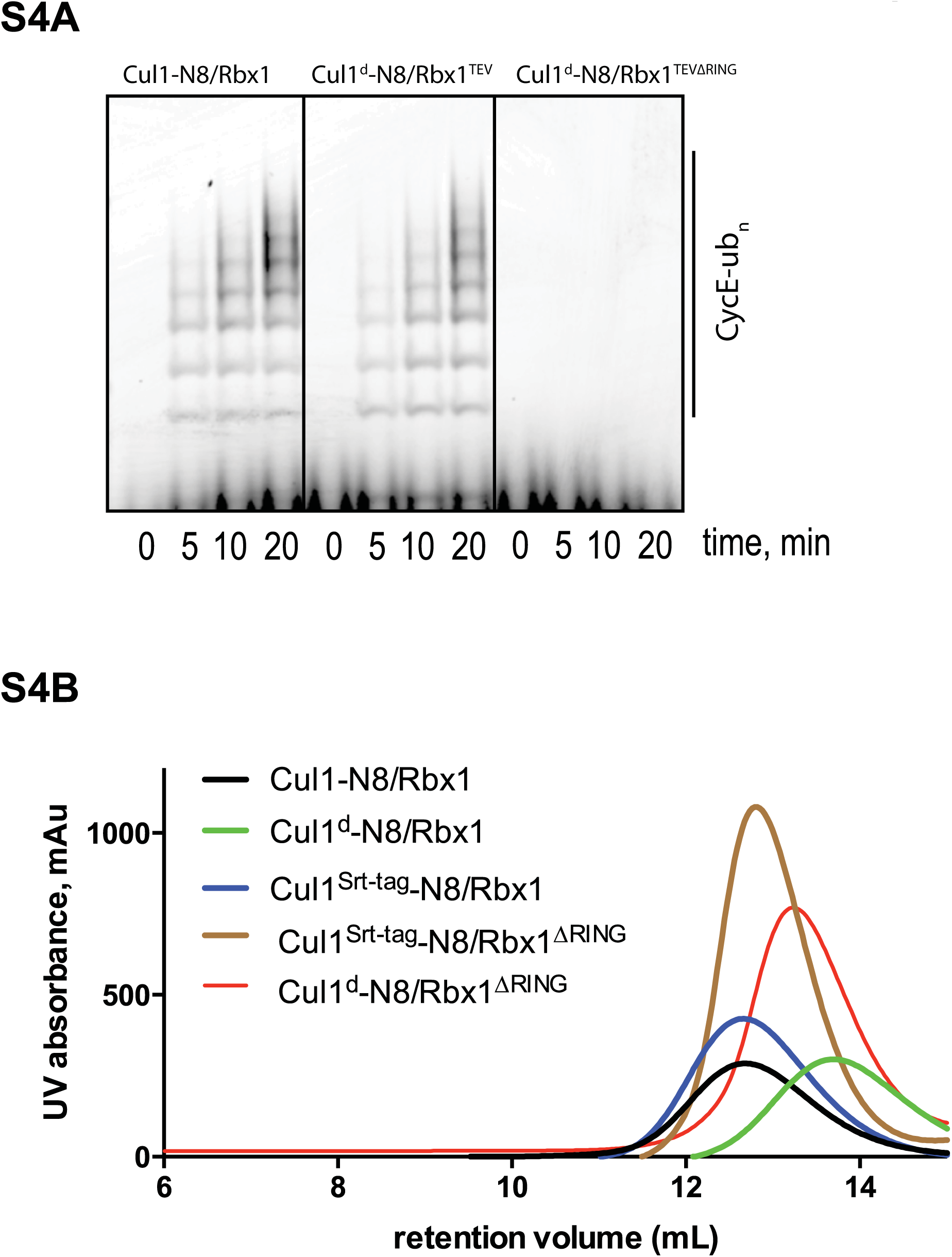
**Biochemical characterization of Cul1/Rbx1^TEVΔRING^ proteins**. (A) Ubiquitination assay using the indicated Cul1-N8/Rbx1 variants (500 nM each) as an E3. Each reaction contained, in addition, 100 nM Ube1, 1000 nM Cdc34b, 750 nM Skp1/Fbxw7 and 4000 nM CyclinE phosphopeptide, labeled with FAM. The samples were incubated at 25°C for the indicated time points, analyzed by SDS PAGE and visualized by excitation at 473 nm. (B) Overlay of Superdex 200 size exclusion profiles of purified Cul1/Rbx1 variants isolated from insect cells.

**Figure 5–figure supplement 1.**
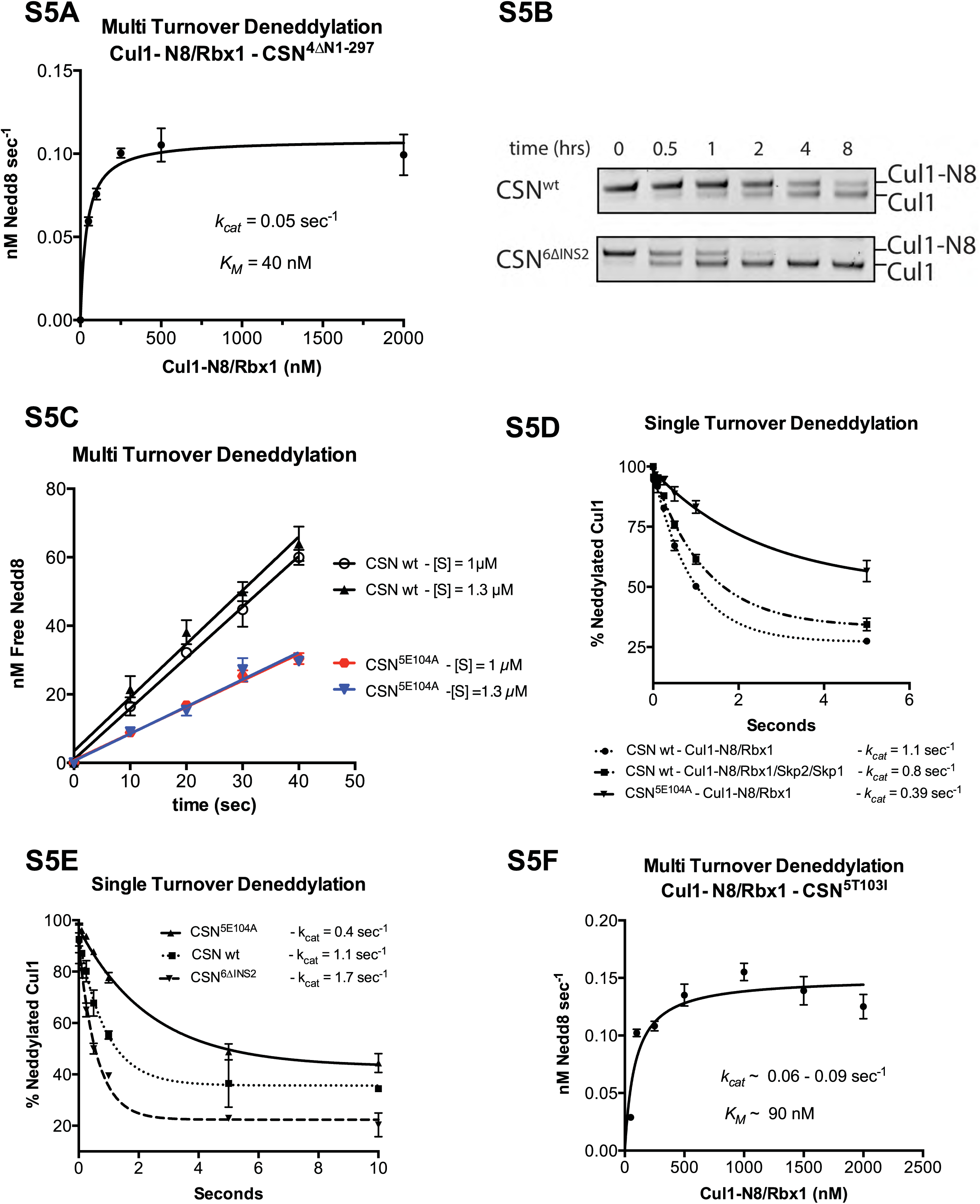
**Kinetic analysis of deneddylation**. (A) Deneddylation reactions were carried out in triplicate with CSN^4Δn^ at varying concentrations of Cul1-[^32^P]N8/Rbx1 substrate and quantified to generate the curve shown. Estimates of *k_cat_* and *K_M_* are indicated. (B) Deneddylation assays of Cul1-N8/Rbx1^ΔRING^ (100 nM), incubated with CSN (200 nM, upper panel) or CSN6^Δins2^ (200 nM, lower panel). Samples were taken at the indicated time points, and visualized by SDS PAGE and Sypro Ruby staining. Note that the ΔRING substrate contained an unreacted Sortase tag at the C-terminus of Cul1 that reduced *k_cat_* by ~4-fold. (C) Multi-turnover deneddylation reactions were carried out with CSN or CSN^5E104A^ and Cul1-[^32^P]N8/Rbx1. Substrate was assayed at 1 and 1.3 µM to confirm that saturation was achieved. (D) Single-turnover deneddylation reactions were carried out with CSN on Cul1-[32P]N8/Rbx1 +/-Skp1/Skp2, and with CSN^5E104A^ on Cul1-[^32^P]N8/Rbx1. (E) Same as panel D except that CSN6^Δins2^ was also evaluated. (F) Multi-turnover deneddylation reactions were carried out in triplicate with CSN5^t103i^ at varying concentrations of Cul1-[^32^P]N8/Rbx1 substrate and quantified to generate the curve shown. Estimates of *k_cat_* and *K_M_* are indicated.

**Figure 6–figure supplement 1.**
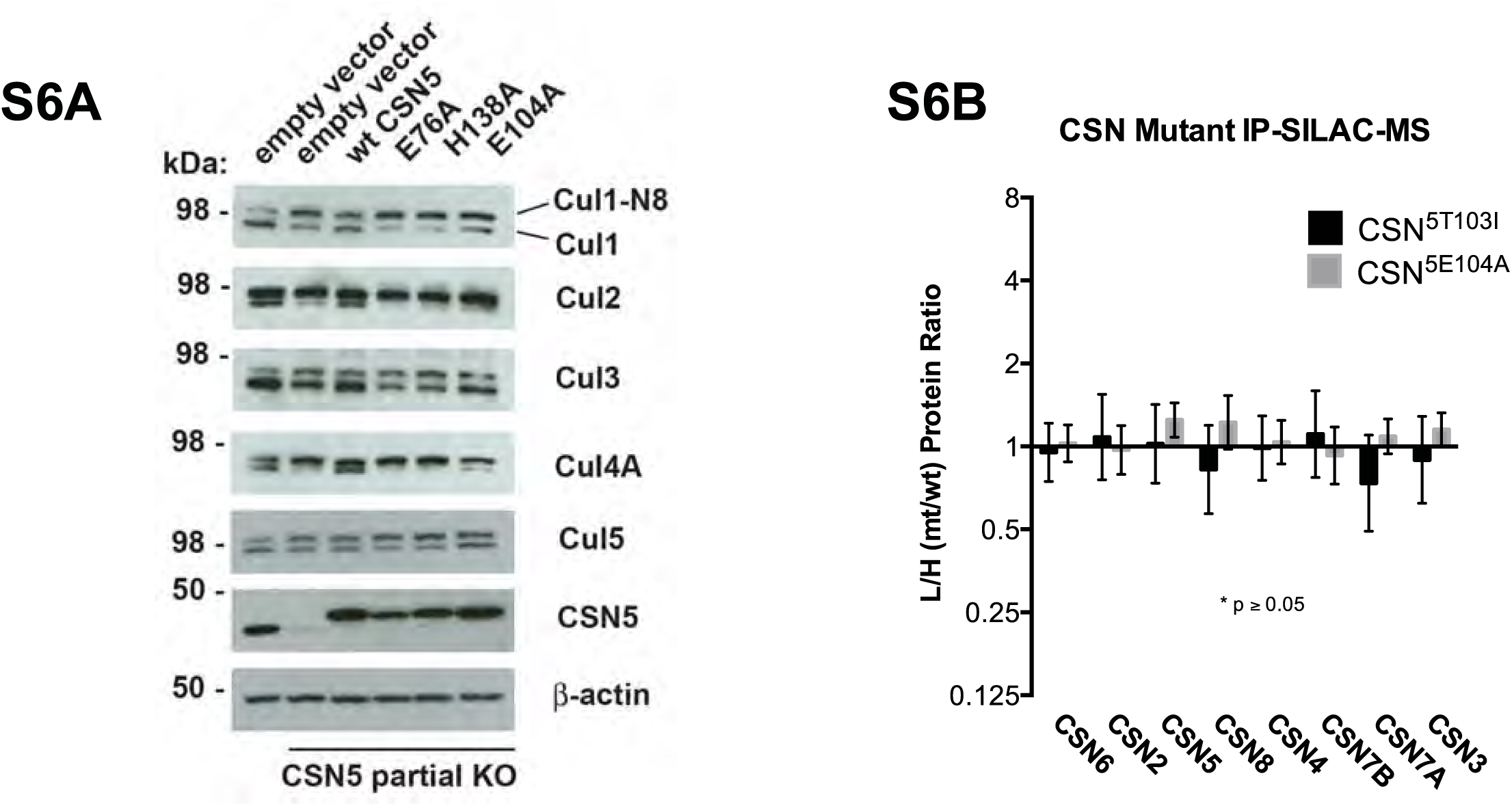
**Time course and titration data for Figure 6A and supplementary immunoblot for Figure 6B**. (A) Same as Figure 6B except that samples were immunoblotted for different cullins. (B) SILAC data for CSN subunits from pull-down analysis shown in Figure 6D.

**Table S1**. **Cross-links within CSN^5H138A^-SCF-N8^Skp2/Cks1^**. “Id” gives the amino acid sequence of the cross-linked peptides and the exact position of the two cross-linked lysine residues is indicated by the numbers of the letters *a* and *b* respectively for the first and second peptide. “**Protein1”** and “**Protein2**” denote the cross-linked protein names and “**Residue 1”** and “**Residue 2”** respectively defines the position of the cross-linked lysine within the sequence of the protein. “**deltaS”** is the delta score of the respective cross-link, which serves as a measure for how close the best assigned hit was scored in regard to the second best. “**Id_Score**” is a weighted sum of four subscores: xcorrc, xcorrx, match-odds and TIC that is used to assess the quality of the composite MS2 spectrum as calculated by *xQuest*. “**FDR**” denotes the false-discovery rate as calculated by *xProphet*. The measured distance in Å is given for all cross-links, which fall within modeled residues.

**Table S2. Cross-links within CSN5^H138A^-SCF^Skp2/Cks1^**

**Table S3. Cross-links within CSN-Cul1/Rbx1**

**Table S4. Cross-links within CSN- SCF-N8^Fbxw7FL^**

**Table S5. Cross-links within CSN- SCF-N8^Fbxw7trunc^**

**Table S6. Cross-links within CSN- Cul3/Rbx1**

**Movie 1**: **Morphing CSN and Cul1-N8/Rbx1 conformational changes, occurring upon binding**. Color code as in Figure 1.

## References

Ambroggio XI, Rees DC, Deshaies RJ 2004. JAMM: A Metalloprotease-Like Zinc Site in the Proteasome and Signalosome. PLoS Biol 2: E2.

Bennett EJ, Rush J, Gygi SP, Harper JW 2010. Dynamics of cullin-RING ubiquitin ligase network revealed by systematic quantitative proteomics. Cell 143: 951–965.

Berger I, Fitzgerald DJ, Richmond TJ 2004. Baculovirus expression system for heterologous multiprotein complexes. Nat Biotechnol 22: 1583–1587.

Birol M, Enchev RI, Padilla A, Stengel F, Aebersold R, Betzi S, Yang Y, Hoh F, Peter M, Dumas C, et al. 2014. Structural and biochemical characterization of the Cop9 signalosome CSN5/CSN6 heterodimer. PLoS One 9: e105688.

Briggs GE, Haldane JB 1925. A Note on the Kinetics of Enzyme Action. Biochem J 19: 338–339.

Cope, GA, Deshaies RJ 2006. Targeted silencing of Jab1/Csn5 in human cells downregulates SCF activity through reduction of F-box protein levels. BMC Biochem 7: 1.

Cope GA, Suh GS, Aravind L, Schwarz SE, Zipursky S., Koonin EV, Deshaies RJ 2002. Role of predicted metalloprotease motif of Jab1/Csn5 in cleavage of Nedd8 from Cul1. Science 298: 608–611.

Cox J, Mann M 2008. MaxQuant enables high peptide identification rates, individualized p.p.b.-range mass accuracies and proteome-wide protein quantification. Nat Biotechnol 26: 1367–1372.

Cox J, Neuhauser N, Michalski A, Scheltema RA, Olsen J., Mann M 2011. Andromeda: a peptide search engine integrated into the MaxQuant environment. J Proteome Res 10: 1794–1805.

den Besten, W, Verma R, Kleiger G, Oania RS, Deshaies RJ 2012. NEDD8 links cullin-RING ubiquitin ligase function to the p97 pathway. Nat Struct Mol Biol 19: 511–516, S511.

Deshaies RJ, Joazeiro CA 2009. RING domain E3 ubiquitin ligases. Annu Rev Biochem 78: 399–434.

Dougherty WG, Cary SM, Parks TD 1989. Molecular genetic analysis of a plant virus polyprotein cleavage site: a model. Virology 171: 356–364.

Duda DM, Borg LA, Scott D., Hunt HW, Hammel M, Schulman BA 2008. Structural insights into NEDD8 activation of cullin-RING ligases: conformational control of conjugation. Cell 134: 995–1006.

Echalier A, Pan Y, Birol M, Tavernier N, Pintard L, Hoh F, Ebel C, Galophe N, Claret FX, Dumas C 2013. Insights into the regulation of the human COP9 signalosome catalytic subunit, CSN5/Jab1. Proc Natl Acad Sci U S A 110: 1273–1278.

Emberley ED, Mosadeghi R, Deshaies RJ 2012. Deconjugation of Nedd8 from Cul1 is directly regulated by Skp1-Fbox and substrate, and CSN inhibits deneddylated SCF by a non-catalytic mechanism. J Biol Chem.

Enchev RI, Schreiber A, Beuron F, Morris EP 2010. Structural insights into the COP9 signalosome and its common architecture with the 26S proteasome lid and eIF3. Structure 18: 518–527.

Enchev RI, Schulman BA, Peter M 2015. Protein neddylation: beyond cullin-RING ligases. Nat Rev Mol Cell Biol 16: 30–44.

Enchev RI, Scott DC, da Fonseca PC, Schreiber A, Monda JK, Schulman B., Peter M, Morris EP 2012. Structural basis for a reciprocal regulation between SCF and CSN. Cell reports 2: 616–627.

Fischer ES, Scrima A, Bohm K, Matsumoto S, Lingaraju GM, Faty M, Yasuda T, Cavadini S, Wakasugi M, Hanaoka F, et al. 2011. The Molecular Basis of CRL4(DDB2/CSA) Ubiquitin Ligase Architecture, Targeting, and Activation. Cell 147: 1024–1039.

Kelley LA, Mezulis S, Yates CM, Wass M., Sternberg MJ 2015. The Phyre2 web portal for protein modeling, prediction and analysis. Nature protocols 10: 845–858.

Leitner A, Walzthoeni T, Aebersold R 2014. Lysine-specific chemical cross-linking of protein complexes and identification of cross-linking sites using LC-MS/MS and the xQuest/xProphet software pipeline. Nature protocols 9: 120–137.

Lingaraju GM, Bunker RD, Cavadini S, Hess D, Hassiepen U, Renatus M, Fisher ES, Thoma NH 2014. Crystal structure of the human COP9 signalosome. Nature 512: 161–165.

Liu J, Furukawa M, Matsumoto T, Xiong Y 2002. NEDD8 modification of CUL1 dissociates p120(CAND1), an inhibitor of CUL1-SKP1 binding and SCF ligases. Mol Cell 10: 1511–1518.

Lyapina S, Cope G, Shevchenko A, Serino G, Tsuge T, Zhou C, Wolf DA, Wei N, Deshaies RJ 2001. Promotion of NEDD-CUL1 conjugate cleavage by COP9 signalosome. Science 292: 1382–1385.

Mindell JA, Grigorieff N 2003. Accurate determination of local defocus and specimen tilt in electron microscopy. J Struct Biol 142: 334–347.

Orthwein 2015. Induction of homologous recombination in G1 cells. Nature in press.

Pettersen, EF, Goddard TD, Huang C., Couch GS, Greenblatt D., Meng EC, Ferrin TE 2004. UCSF Chimera-a visualization system for exploratory research and analysis. J Comput Chem 25: 1605–1612.

Pierce NW, Lee JE, Liu X, Sweredoski MJ, Graham R., Larimore EA, Rome M, Zheng N, Clurman BE, Hess S, et al. 2013. Cand1 promotes assembly of new SCF complexes through dynamic exchange of F box proteins. Cell 153: 206–215.

Politis A, Stengel F, Hall Z, Hernandez H, Leitner A, Walzthoeni T, Robinson CV, Aebersold R 2014. A mass spectrometry-based hybrid method for structural modeling of protein complexes. Nat Methods 11: 403–406.

Ritchie ME, Phipson B, Wu D, Hu Y, Law CW, Shi W, Smyth GK 2015. limma powers differential expression analyses for RNA-sequencing and microarray studies. Nucleic Acids Res 43: e47.

Rosenthal PB, Henderson R 2003. Optimal determination of particle orientation, absolute hand, and contrast loss in single-particle electron cryomicroscopy. J Mol Biol 333: 721–745.

Saha A, Deshaies RJ 2008. Multimodal activation of ubiquitin ligase SCF by Nedd8 conjugation. Mol Cell 32: 21–31.

Sato Y, Yoshikawa A, Yamagata A, Mimura H, Yamashita M, Ookata K, Nureki O, Iwai K, Komada M, Fukai S 2008. Structural basis for specific cleavage of Lys 63-linked polyubiquitin chains. Nature 455: 358–362.

Scheres, SH 2012. RELION: implementation of a Bayesian approach to cryo-EM structure determination. J Struct Biol 180: 519–530.

Scheres SH, Chen S 2012. Prevention of overfitting in cryo-EM structure determination. Nat Methods 9: 853–854.

Schmidt MW, McQuary PR, Wee S, Hofmann K, Wolf DA 2009. F-box-directed CRL complex assembly and regulation by the CSN and CAND1. Mol Cell 35: 586–597.

Shalem O, Sanjana NE, Hartenian E, Shi X, Scott DA, Mikkelsen T., Heckl D, Ebert BL, Root D., Doench JG, et al. 2014a. Genome-scale CRISPR-Cas9 knockout screening in human cells. Science 343: 84–87.

Shalem O, Sanjana NE, Hartenian E, Shi X, Scott DA, Mikkelsen T., Heckl D, Ebert BL, Root D., Doench JG, et al. 2014b. Genome-scale CRISPR-Cas9 knockout screening in human cells. Science 343: 3.

Skaar JR, Pagan JK, Pagano M 2013. Mechanisms and function of substrate recruitment by F-box proteins. Nat Rev Mol Cell Biol 14: 369–381.

Smyth, GK (2005). Limma: linear models for microarray data. In Bioinformatics and computational biology solutions using R and Bioconductor (New York: Springer), pp. 397–420.

Soucy TA, Smith PG, Milhollen M., Berger AJ, Gavin J., Adhikari S, Brownell JE, Burke K., Cardin DP, Cullis CA 2009. An inhibitor of NEDD8-activating enzyme as a novel approach to treat cancer. Nature 458: 732–736.

Suh GS, Poeck B, Chouard T, Oron E, Segal D, Chamovitz DA, Zipursky SL 2002. Drosophila JAB1/CSN5 acts in photoreceptor cells to induce glial cells. Neuron 33: 35–46.

Theile CS, Witte MD, Blom A., Kundrat L, Ploegh HL, Guimaraes CP 2013. Site-specific N-terminal labeling of proteins using sortase-mediated reactions. Nature protocols 8: 1800–1807.

Tran HJ, Allen MD, Lowe J, Bycroft M 2003. Structure of the Jab1/MPN domain and its implications for proteasome function. Biochemistry 42: 11460–11465.

Wee S, Geyer RK, Toda T, Wolf DA 2005. CSN facilitates Cullin-RING ubiquitin ligase function by counteracting autocatalytic adapter instability. Nat Cell Biol 7: 387–391.

Wu S, Zhu W, Nhan T, Toth JI, Petroski M., Wolf DA 2013. CAND1 controls in vivo dynamics of the cullin 1-RING ubiquitin ligase repertoire. Nature communications 4: 1642.

Yamoah K, Oashi T, Sarikas A, Gazdoiu S, Osman R, Pan ZQ 2008. Autoinhibitory regulation of SCF-mediated ubiquitination by human cullin 1’s C-terminal tail. Proc Natl Acad Sci U S A 105: 12230–12235.

Zemla A, Thomas Y, Kedziora S, Knebel A, Wood NT, Rabut G, Kurz T 2013. CSN- and CAND1-dependent remodelling of the budding yeast SCF complex. Nature communications 4: 1641.

Zheng N, Schulman BA, Song L, Miller JJ, Jeffrey P., Wang P, Chu C, Koepp DM, Elledge S., Pagano M, et al. 2002. Structure of the Cul1-Rbx1-Skp1-F boxSkp2 SCF ubiquitin ligase complex. Nature 416: 703–709.

